# Variant-to-function mapping of late-onset Alzheimer’s disease GWAS signals in human microglial cell models implicates *RTFDC1* at the *CASS4* locus

**DOI:** 10.1101/2024.08.22.609230

**Authors:** Elizabeth A. Burton, Mariana Argenziano, Kieona Cook, Molly Ridler, Sumei Lu, Chun Su, Elisabetta Manduchi, Sheridan H. Littleton, Michelle E. Leonard, Kenyaita M. Hodge, Li-San Wang, Gerard D. Schellenberg, Matthew E. Johnson, Matthew C. Pahl, James A. Pippin, Andrew D. Wells, Stewart A. Anderson, Christopher D. Brown, Struan F.A. Grant, Alessandra Chesi

## Abstract

Late-onset Alzheimer’s disease (LOAD) research has principally focused on neurons over the years due to their known role in the production of amyloid beta plaques and neurofibrillary tangles. In contrast, recent genomic studies of LOAD have implicated microglia as culprits of the prolonged inflammation exacerbating the neurodegeneration observed in patient brains. Indeed, recent LOAD genome-wide association studies (GWAS) have reported multiple loci near genes related to microglial function, including *TREM2*, *ABI3*, and *CR1*. However, GWAS alone cannot pinpoint underlying causal variants or effector genes at such loci, as most signals reside in non-coding regions of the genome and could presumably confer their influence frequently via long-range regulatory interactions. We elected to carry out a combination of ATAC-seq and high-resolution promoter-focused Capture-C in two human microglial cell models (iPSC-derived microglia and HMC3) in order to physically map interactions between LOAD GWAS-implicated candidate causal variants and their corresponding putative effector genes. Notably, we observed consistent evidence that rs6024870 at the GWAS *CASS4* locus contacted the promoter of nearby gene, *RTFDC1*. We subsequently observed a directionallly consistent decrease in *RTFDC1* expression with the the protective minor A allele of rs6024870 via both luciferase assays in HMC3 cells and expression studies in primary human microglia. Through CRISPR-Cas9-mediated deletion of the putative regulatory region harboring rs6024870 in HMC3 cells, we observed increased pro-inflammatory cytokine secretion and decreased DNA double strand break repair related, at least in part, to *RTFDC1* expression levels. Our variant-to-function approach therefore reveals that the rs6024870-harboring regulatory element at the LOAD ‘*CASS4’* GWAS locus influences both microglial inflammatory capacity and DNA damage resolution, along with cumulative evidence implicating *RTFDC1* as a novel candidate effector gene.

## INTRODUCTION

Late-onset Alzheimer’s disease (LOAD) is the most common neurodegenerative disease presenting over the age of 65, affecting nearly 6 million American adults^1^. Despite being the seventh leading cause of death in the United States, there are no effective therapies that can alter disease progression. LOAD risk increases exponentially with age, with 3-5% presenting with LOAD at 65-69 years old but 30-40% by 80^2^.

LOAD is pathologically characterized by the accumulation of extracellular Aβ_1-42_ plaques and intraneuronal neurofibrillary tangles consisting of hyperphosphorylated tau protein, which are widely proposed to progressively lead to neuronal cell death. As a consequence, the majority of hypothesis-driven work in the LOAD field has focused on characterizing the impact of Aβ_1-42_ plaques and neurofibrillary tangles on neurons. However, recent genetic findings are yielding fresh insights.

Population studies have shown that approximately 60-80% of LOAD risk is explained by heritable factors^3^, suggesting a strong polygenic component. The most recent large-scale genome wide association study (GWAS), a method which which offers unbiased identification of susceptibility loci, identified over 70 independent signals associated with LOAD^4^, including variants near the *MAPT (*encoding tau) and *APP* (encoding the amyloid precursor protein) genes. However, a meta-analysis of GWAS colocalized genes showed strong enrichment in genes active in immune-related tissues, including the microglia^5^, the resident macrophages of the central nervous system (CNS).

Indeed, microglia have received increased attention as a key cell type in the pathogenesis of LOAD. In response to Aβ_1-42_ production, microglia activate through a gene regulatory cascade and migrate to the site of plaque accumulation, where they then break down and phagocytose the amyloid plaques,^6,7^ resulting in the release of pro-inflammatory cytokines^8,9^. Persistent production of these cytokines reduces the ability of the microglia to clear Aβ_1-42_ within a negative feedback loop, which also increases production of hyperphosphorylated tau, driving heightened neuroinflammation which leads ultimately to neuronal death^10,11^. Additionally, linkage disequilibrium (LD) score regression of LOAD GWAS-associated variants showed greatest enrichment in microglia compared to all other brain cell types, further suggesting that this cell type plays a central role in LOAD pathogenesis^12^. Consistent with these enrichment analyses, both rare and common variants have been identified in core microglial genes, such as *TREM2*^13,14^. Taken together, these observations implicate microglia as a major contributing cell type to LOAD pathology.

A limitation of GWAS is that the approach only reports the sentinel single nucleotide polymorphism (SNP) at a given locus, which serves as a tag for all other variants in LD; as such, among this entire LD series of proxy SNPs, any one (or more) could be causal^15^. Furthermore, the majority of these sentinel SNPs and their associated proxies lie within non-coding regions of the genome, and may not necessarily influence the expression of the nearest gene. Instead, these variants likely modulate the activity of distal regulatory elements, such as enhancers, which in turn regulate distal gene expression^16^, which is frequently not the nearest gene. One such example comes from the strongest associated obesity GWAS signal that lies within an intron of the *FTO* gene, but principally influences the expression of distal genes *IRX3* and *IRX5*^17–19^. Additionally, we identified *ING3* and *EPDR1* as the long-range effector genes regulated by osteoblast enhancers harboring bone mineral density GWAS SNPs at the *WNT16-CPED1* and *STARD3NL* loci, respectively^20^.

Given that many regulatory elements have the capacity to influence gene expression via long-range interactions across relatively large genomic distances, and that many of these elements function in a cell-type specific manner^21^, identification of active GWAS-implicated enhancers in microglial cells should drive cell-type specific insights in to the genetic etiology of LOAD and yield more therapeutically relevant targets. Indeed, the causal SNP at the LOAD GWAS-implicated *BIN1* locus resides within an enhancer that is only active in the microglia, and not other CNS cell types^22^. As such, we elected to identify chromatin contacts between LOAD GWAS-associated variants^4,23^ and their corresponding effector genes via a “variant-to-gene mapping” strategy utilizing a combination of a high-resolution chromatin conformation-based Capture-C assay coupled with chromatin accessibility assessed by ATAC-seq^24–30^.

To this end, we performed our variant-to-gene mapping approach in two microglial cell models: induced pluripotent stem cell (iPSC)-derived human microglia (iMg) and the human microglial clone 3 (HMC3) immortalized cell line. Our results consistently pointed to a contact at the LOAD GWAS-implicated *CASS4* locus between rs6024870 and the promoter of neighboring gene *RTFDC1.* Given the relative tractability of the HMC3 model, we pursued functional validation in this cell line via CRISPR-mediated precise deletion of an approximately 400bp region harboring rs6024870.. Altogether, we present variant-to-function evidence that the genomic region harboring rs6024870 acts as a microglial enhancer and influences LOAD risk through regulation of both inflammatory and DNA damage related pathways.

## RESULTS

### Variant-to-gene mapping in two human microglia models implicates putative causal variants and corresponding effector genes at LOAD GWAS loci

To implicate candidate causal genetic variants at LOAD loci and to map them to their corresponding putative effector genes, we leveraged the signals reported in the two most recently published LOAD GWAS meta-analyses^4,31^. We extracted all 111 sentinel SNPs from these studies (83 from Bellenguez *et al.*; 38 from Wightman *et al.*; 10 index SNPs were in common). Ten of these sentinel SNPs were in high LD (r^2^>0.7), so we retained one representative SNP from each pair, resulting in 101 SNPs under consideration in total. For each sentinel SNP, we obtained all proxy SNPs (r^2^>0.7) using the European panel of the 1000 genomes phase 3 v.5 available from SNiPA^32^.^32^. This LD expansion process yielded 2,835 candidate LOAD-associated ‘proxy’ SNPs.

To facilitate physical fine-mapping of these candidate variants and to subsequently implicate their corresponding effector genes, we elected to use two different cell models, the human microglial cell line HMC3, and iPSC-derived microglia (iMg)^33^. While HMC3 is an immortalized cell line, it is positive for the microglial/macrophage marker IBA1 and the endotoxin receptor CD14, and negative for the astrocyte marker GFAP^34^. iMg, on the other hand, are derived from hematopoietic precursor cells (HPCs), which are the normal precursor cells of adult macrophages^35^. While adult microglia are autonomously maintained in the CNS independent of HPCs once mature^36^, derivation of iMg from HPCs recapitulates the early yolk sac HPCs that give rise to embryonic microglia^37–39^, and thus represents a useful human microglial model.

To first confirm that these lines were appropriate models to conduct our variant-to-gene mapping campaign, we performed RNA-seq (**Suppl. Table 1**) and compared the transcriptomes to published human brain-isolated microglia (*ex-vivo* or from post-mortem brains^22,40^) plus other cell types from the GTEx Analysis RNA-seq databasel (v. 7)^41^ and monocyte and macrophage datasets^42,43^. The transcriptome of iMg closely resembled that of *ex-vivo* microglia, and more so than other related innate immune cell types such as macrophages and monocytes (**Figure 1 A, B, C**). HMC3 also clustered close to other microglial-like cell types, supporting the utility of this cell type as a tractable microglial model in our functional follow-up experiments.

**Figure 1:**
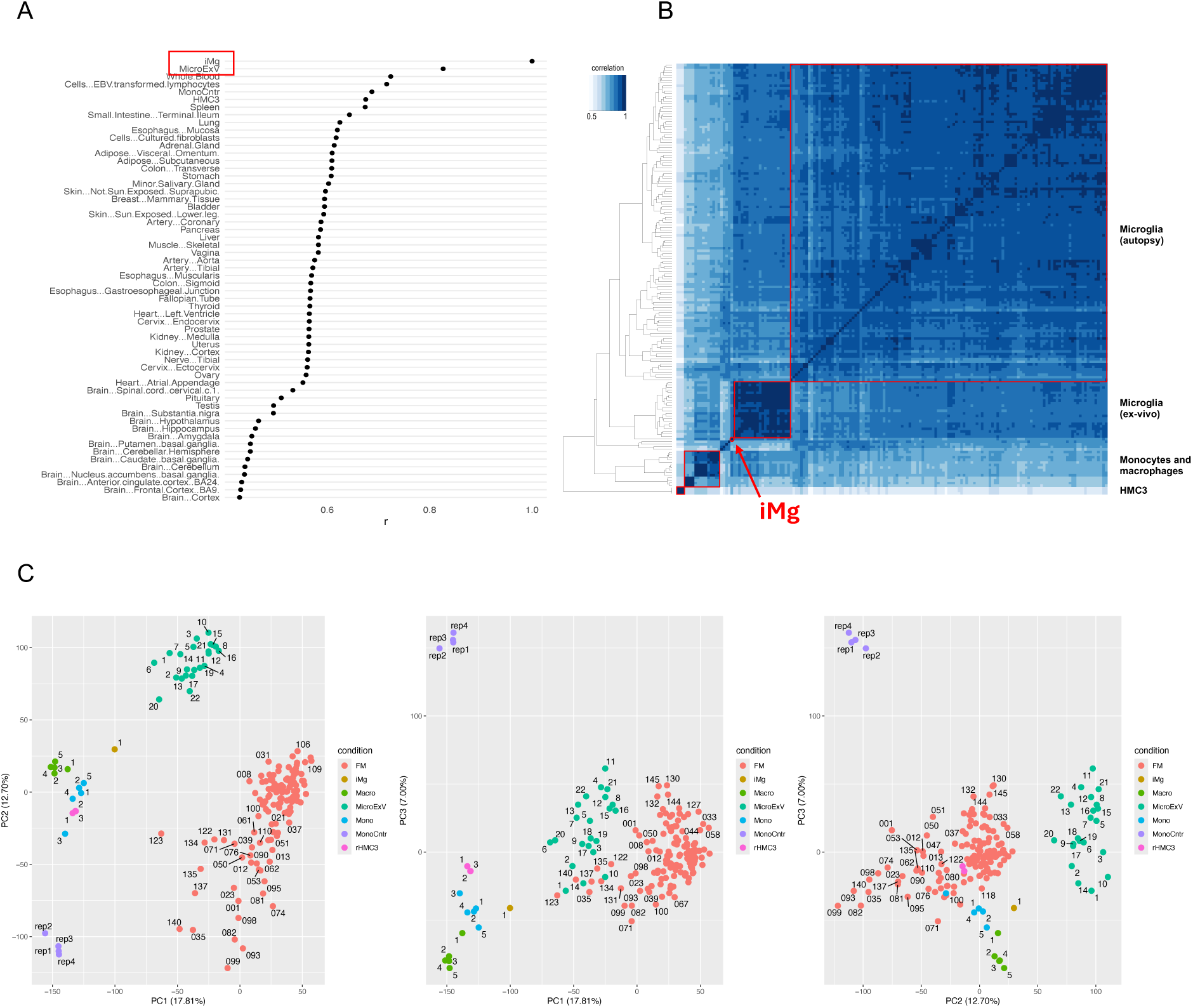
iPSC-derived microglia (iMg) and HMC3 transcriptomes from this study show high correlation with established *in vivo* human microglia datasets. **(A)** Comparison of iMg expression profile to tissues in the GTEx database. R scores are the Spearman correlation coefficient of 16,651 genes expressed in both datasets. Median expression values were downloaded from GTEx (v7); MicroExV from Nott *et al.*^22^; Mono Cntr from Pahl et al.^43^; iMg and HMC3 are from this study. **(B)** Heatmap of the correlation of gene expression between microglia-related cell types clustered by distance. Samples shown are: microglia from autopsies of 127 individuals from the FreshMicro study^40^, ex-vivo microglia from biopsies of 22 individuals^22^, monocytes and macrophages^42,89^, monocytes^43^, and iMg and HMC3 from this study. **(C)** Principal component analysis of the same datasets as in B.

To physically fine-map the LOAD associated proxy SNPs using cell-specific open chromatin maps we performed ATAC-seq, generating three replicate libraries from both cell types according to standard protocols, and sequenced them on the Illumina platform. Peaks were called with the ENCODE ATAC-seq pipeline (https://www.encodeproject.org/atac-seq/) using the optimal peaks from the IDR procedure, which resulted in 145,009 and 55,094 open chromatin peaks for iMg and HMC3, respectively (**Suppl. Table 2**). We intersected the genomic positions of the proxy LOAD SNPs with the open ATAC-seq peaks to obtain 156 “open” prioritized proxy SNPs in iMg and 29 in HMC3 (42 were shared between the two cell types). Thus, filtering candidate proxies using open chromatin peaks from ATAC-seq effectively reduced the number of candidates by a factor of ∼20-100.

Next, to connect LOAD-associated SNPs to potential effector genes, we performed high-resolution promoter-focused Capture-C in both iMg and HMC3 cells. We generated three replicate libraries following our previously published protocol^20^ and sequenced them on the Illumina platform. We pre-processed the reads using HICUP^44^. Libraries from each cell type yielded high coverage (an average of ∼2.6 billion reads per library) with >70% capture efficiency (% of unique valid reads captured; **Suppl. Table 3**). We called chromatin interactions using CHiCAGO^45^ with standard parameters at two resolutions, i.e. one and four DpnII fragments to boost sensitivity for both short-range and long-range interactions^46^.

Overall, we identified a total of 362,773 *cis* interactions in iMg cells (∼5% being bait-to-bait), with a median interaction distance of 61.8 kb (**Suppl. Table 4)**. Most of the non-baited promoter-interacting regions (PIRs) had contacts with a single baited region (91%), while only 0.2% contacted more than four. PIRs were significantly enriched for open chromatin regions detected in our ATAC-seq experiments and for active chromatin marks from *ex-vivo* microglia^22^, suggesting a potential regulatory role (**Suppl. Fig 1**).

We then performed partitioned heritability LD Score regression^47^ to identify enrichment of LOAD GWAS polygenic heritability among the Capture-C-prioritized regions identified from iMg, using GWAS summary statistics for schizophrenia as a control^48^. The LOAD GWAS signals were significantly enriched in iMg (*P*=0.04, enrichment 47.2), while schizophrenia signals were not (*P*=0.81, enrichment 1.5), which is in line with expectation given that schizophrenia loci are widely hypothesized to confer their effects principally via neuronal cell types.

We obtained similar results for the HMC3 cells, identifying 525,306 *cis* interactions (13% bait-to-bait) with a median interaction distance of 49.4 kb, 74% of the PIRs interacting with a single bait fragment, and 3.7% with more than four (**Suppl. Table 4)**. PIRs in HMC3 were similarly enriched for active chromatin markers (**Suppl. Fig. 1**). All Capture-C interactions for the two cell types are reported in **Suppl. Table 4.**

We went on to map all LOAD GWAS proxies residing in open chromatin to their putative effector genes. In the two microglial cell models, 16 LOAD loci revealed at least one or more proxy SNPs in open chromatin (and not residing in a baited promoter region) interacting with an open gene promoter. A total of 32 open baited regions corresponding to 34 gene promoters (25 coding and 9 non-coding) contacted 34 open chromatin regions harboring one or more LOAD proxy SNPs (34 total) through 36 distinct contacts (**Table 1**). Of these contacts, 7 (19%) were to the nearest gene, 8 (22%) to both nearest and more distal genes, and 21 (58%) to distal genes only. 26 (79%) of the SNPs contacted one effector gene and 7 (21%) more than one. Five genes (*SLC8B1*, *RTFDC1*, *MS4A6A*, *BLNK*, and *PTK2B*) were contacted by more than one LOAD SNP-containing region (multiple putative enhancers).

**Table 1:**
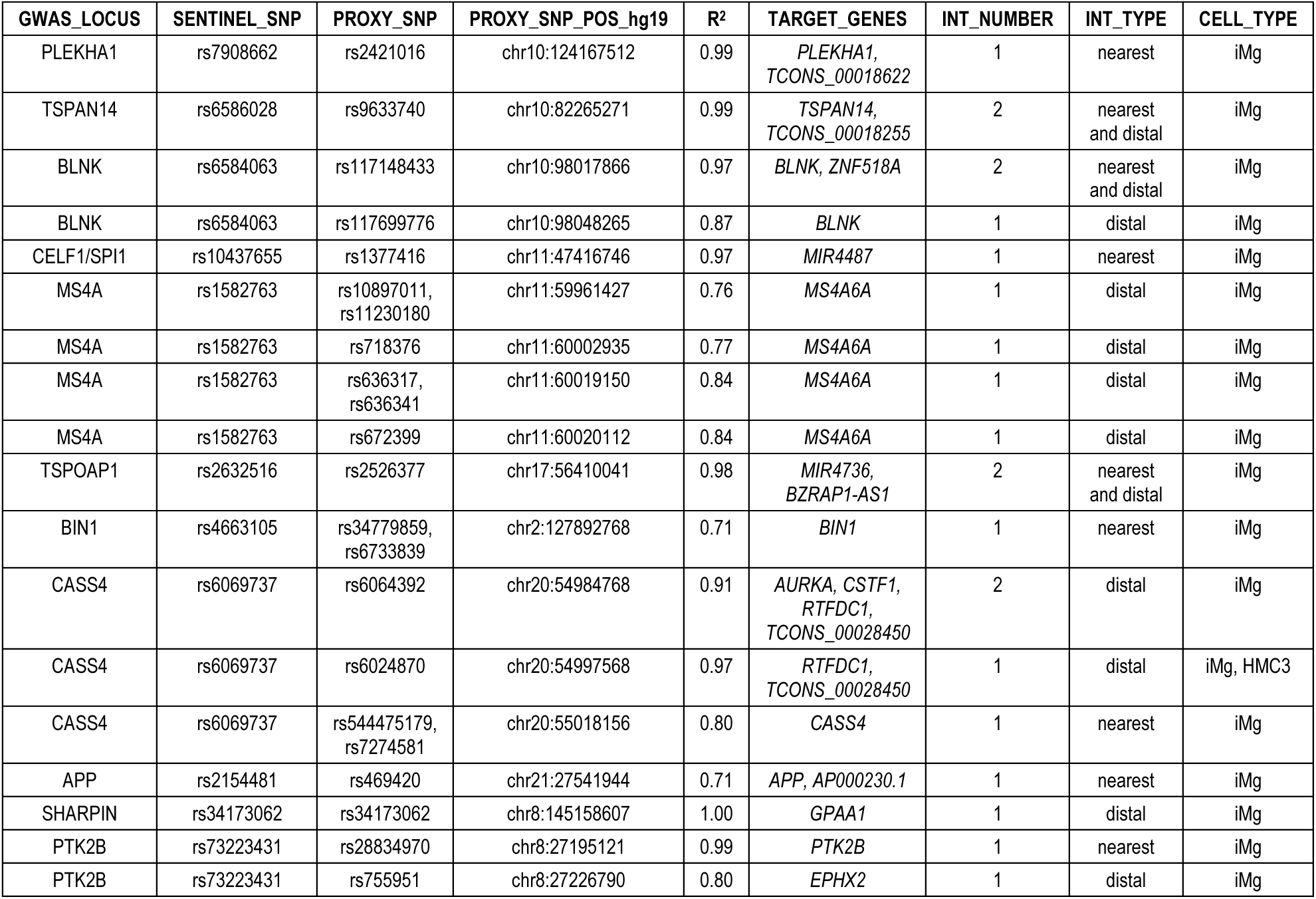
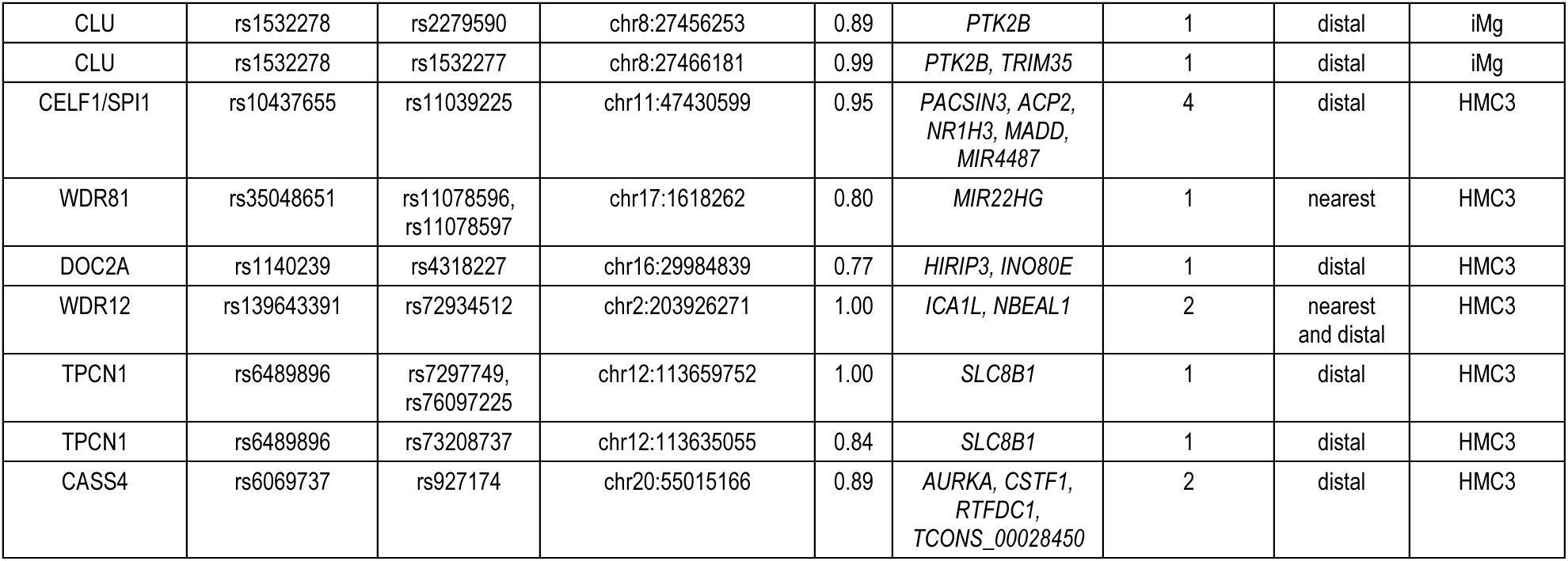
Implicated target genes at 16 LOAD GWAS loci in microglial cells. For each GWAS locus, proxy SNPs in open chromatin contacting an open gene promoter are reported, together with their sentinel SNPs, r^2^, target genes, and the number and type of interaction. Looping interactions and open chromatin maps were derived from promoter-focused Capture-C and ATAC-seq experiments in iMg and HMC3 cells.

| GWAS_LOCUS | SENTINEL_SNP | PROXY_SNP | PROXY_SNP_POS_hg19 | R<sup>2</sup> | TARGET_GENES | INT_NUMBER | INT_TYPE | CELL_TYPE |
| --- | --- | --- | --- | --- | --- | --- | --- | --- |
| PLEKHA1 | rs7908662 | rs2421016 | chr10:124167512 | 0.99 | *PLEKHA1*,<br>*TCONS_00018622* | 1 | nearest | iMg |
| TSPAN14 | rs6586028 | rs9633740 | chr10:82265271 | 0.99 | *TSPAN14*,<br>*TCONS_00018255* | 2 | nearest<br>and distal | iMg |
| BLNK | rs6584063 | rs117148433 | chr10:98017866 | 0.97 | *BLNK*, *ZNF518A* | 2 | nearest<br>and distal | iMg |
| BLNK | rs6584063 | rs117699776 | chr10:98048265 | 0.87 | *BLNK* | 1 | distal | iMg |
| CELF1/SPI1 | rs10437655 | rs1377416 | chr11:47416746 | 0.97 | *MIR4487* | 1 | nearest | iMg |
| MS4A | rs1582763 | rs10897011,<br>rs11230180 | chr11:59961427 | 0.76 | *MS4A6A* | 1 | distal | iMg |
| MS4A | rs1582763 | rs718376 | chr11:60002935 | 0.77 | *MS4A6A* | 1 | distal | iMg |
| MS4A | rs1582763 | rs636317,<br>rs636341 | chr11:60019150 | 0.84 | *MS4A6A* | 1 | distal | iMg |
| MS4A | rs1582763 | rs672399 | chr11:60020112 | 0.84 | *MS4A6A* | 1 | distal | iMg |
| TSPOAP1 | rs2632516 | rs2526377 | chr17:56410041 | 0.98 | *MIR4736*,<br>*BZRAP1-AS1* | 2 | nearest<br>and distal | iMg |
| BIN1 | rs4663105 | rs34779859,<br>rs6733839 | chr2:127892768 | 0.71 | *BIN1* | 1 | nearest | iMg |
| CASS4 | rs6069737 | rs6064392 | chr20:54984768 | 0.91 | *AURKA*, *CSTF1*,<br>*RTFDC1*,<br>*TCONS_00028450* | 2 | distal | iMg |
| CASS4 | rs6069737 | rs6024870 | chr20:54997568 | 0.97 | *RTFDC1*,<br>*TCONS_00028450* | 1 | distal | iMg, HMC3 |
| CASS4 | rs6069737 | rs544475179,<br>rs7274581 | chr20:55018156 | 0.80 | *CASS4* | 1 | nearest | iMg |
| APP | rs2154481 | rs469420 | chr21:27541944 | 0.71 | *APP*, *AP000230.1* | 1 | nearest | iMg |
| SHARPIN | rs34173062 | rs34173062 | chr8:145158607 | 1.00 | *GPAA1* | 1 | distal | iMg |
| PTK2B | rs73223431 | rs28834970 | chr8:27195121 | 0.99 | *PTK2B* | 1 | nearest | iMg |
| PTK2B | rs73223431 | rs755951 | chr8:27226790 | 0.80 | *EPHX2* | 1 | distal | iMg |
| CLU | rs1532278 | rs2279590 | chr8:27456253 | 0.89 | *PTK2B* | 1 | distal | iMg |
| CLU | rs1532278 | rs1532277 | chr8:27466181 | 0.99 | *PTK2B, TRIM35* | 1 | distal | iMg |
| CELF1/SPI1 | rs10437655 | rs11039225 | chr11:47430599 | 0.95 | *PACSIN3, ACP2, NR1H3, MADD, MIR4487* | 4 | distal | HMC3 |
| WDR81 | rs35048651 | rs11078596, rs11078597 | chr17:1618262 | 0.80 | *MIR22HG* | 1 | nearest | HMC3 |
| DOC2A | rs1140239 | rs4318227 | chr16:29984839 | 0.77 | *HIRIP3, INO80E* | 1 | distal | HMC3 |
| WDR12 | rs139643391 | rs72934512 | chr2:203926271 | 1.00 | *ICA1L, NBEAL1* | 2 | nearest and distal | HMC3 |
| TPCN1 | rs6489896 | rs7297749, rs76097225 | chr12:113659752 | 1.00 | *SLC8B1* | 1 | distal | HMC3 |
| TPCN1 | rs6489896 | rs73208737 | chr12:113635055 | 0.84 | *SLC8B1* | 1 | distal | HMC3 |
| CASS4 | rs6069737 | rs927174 | chr20:55015166 | 0.89 | *AURKA, CSTF1, RTFDC1, TCONS_00028450* | 2 | distal | HMC3 |

### LOAD risk SNP rs6024870 at the *CASS4* locus contacts the promoter of *RTFDC1*

We next sought to functionally validate a key proxy variant and its associated regulatory element from our variant-to-gene mapping effort using the tractable HMC3 cell model. We limited our candidate regulatory element/variant search to those loci that yielded consistent observation in both our HMC3 and iMg datasets. As a consequence, we elected to characterize the putative regulatory element harboring the GWAS-implicated microglial risk proxy variant, rs6024870, at *CASS4* (**Table 1**), a key LOAD locus uncharacterized in previous studies.

rs6024870 lies within the first intron of *CASS4*. We prioritized this variant as it is in strong LD with the sentinel SNP at this locus (rs6069737, r^2^ = 0.97), and lies within open chromatin in both of our microglial cell models, as well as in *ex vivo* human microglia ^22^. In the GWAS, both the minor alleles of the sentinel SNP (T) and rs6024870 (A) are protective against LOAD. Notably, in both HMC3 cells and iMg, this variant lies within a region of open chromatin. To further examine the epigenetic signature of the region harboring this variant, we queried publicly available ChIP-seq datasets of histone marks indicative of regulatory element activity from four major brain cell types (neurons, astrocytes, microglia, and oligodendrocytes) isolated from resected human cortical brain tissue^22^. Of the four cell types, H3K27ac was only enriched in microglia, suggesting that this region acts specifically as an enhancer in microglia. Additionally, rs6024870 is located 19bp upstream of an ENCODE candidate *cis*-regulatory element (ID E2123559)^49^, further supporting its role as a microglia-specific enhancer. Despite residing within the first intron of *CASS4*, our Capture-C data in both HMC3 cells and iMg show a physical contact between the region harboring rs6024870 and the promoter of the neighboring gene, *RTFDC1* (**Figure 2A**).

**Figure 2:**
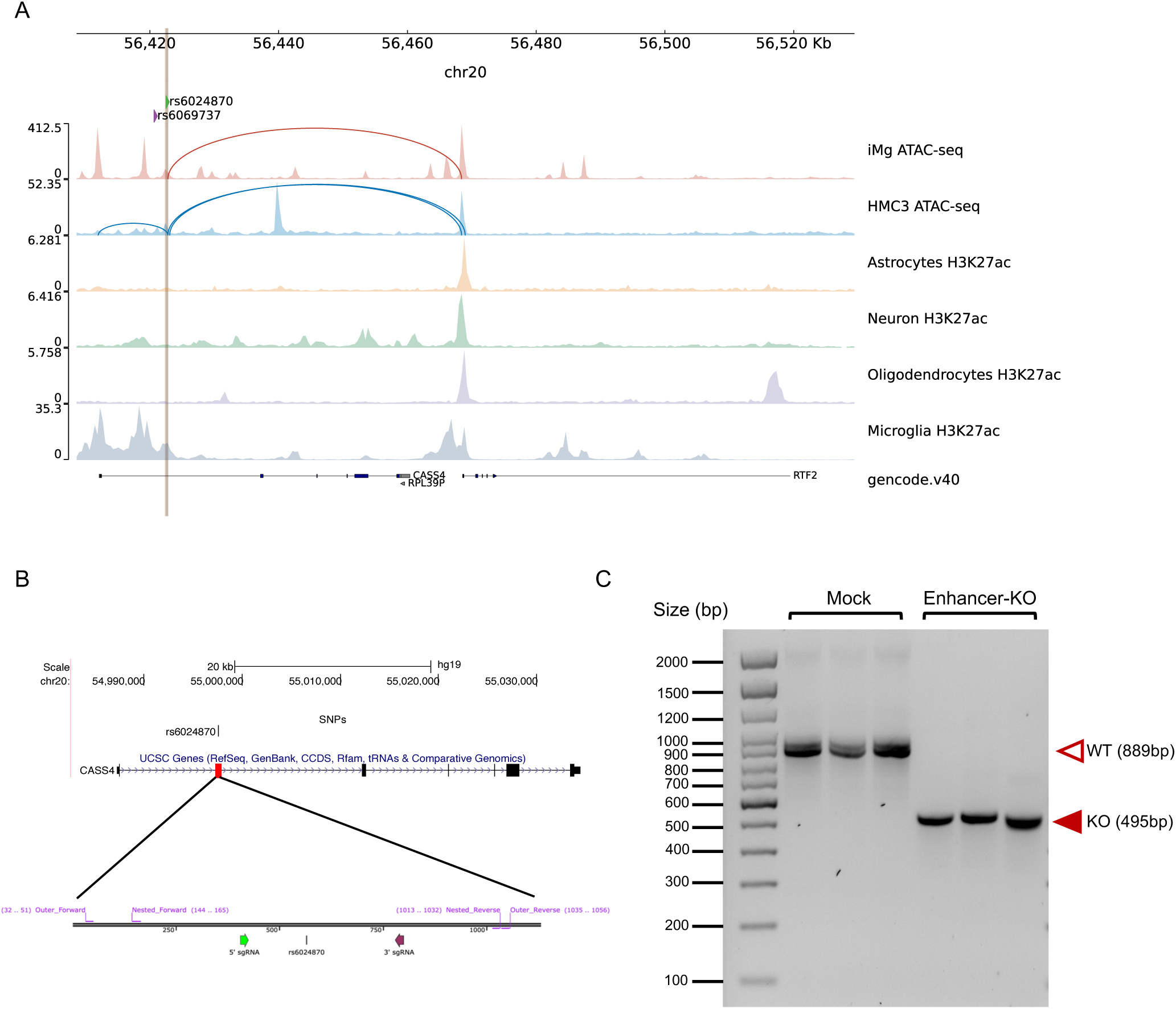
CRISPR-Cas9-mediated deletion of rs6024870-harboring regulatory element in HMC3 Cells. **(A)** Chromatin accessibility peaks from ATAC-seq analysis human microglial cells models iMg cells (red) and HMC3 (blue) at the LOAD *CASS4* GWAS locus. Loops represent physical contacts between regions of DNA as assayed by promoter-focused Capture-C in the respective cell types. Below ATAC-seq peaks, H3K27ac ChIP-seq peaks from Nott *et al*.^22^ for astrocytes (yellow), neurons (green), oligodendrocytes (purple), and microglia (gray). Green arrow and brown line represent the position of rs6024870, while the purple arrow represents the position of the sentinel variant rs6069737. **(B)** Schematic of the CRISPR-Cas9 rs6024870-enhancer harboring regulatory element (enhancer-KO) deletion. Top: UCSC genome browser view of *CASS4* with the deletion region highlighted by the red box. Bottom: SnapGene schematic of sgRNAs (5’ sgRNA: purple arrow; 3’ sgRNA: green arrow) and genotyping PCR primers utilized to confirm successful deletion. **(C)** DNA gel confirming the genotype of 3 mock clones and 3 enhancer-KO clones. Each lane represents a PCR amplification of the deletion region from genomic DNA from an individual clone.

### CRISPR-Cas9 mediated deletion of the rs6024870-harboring enhancer in HMC3

To further characterize the functional effects of the putative enhancer harboring rs6024870, we performed CRISPR-Cas9-mediated genome editing of the rs6024870-harboring enhancer in the HMC3 cell line (**Figure 2B**). While *CASS4* has been previously shown to be associated with LOAD risk and tau toxicity by association studies^50,51^, the mechanism by which *RTFDC1* potentially modulates the pathogenesis of LOAD is unclear. To characterize the loss-of-function effect of this enhancer region, we generated three clones in the HMC3 cell line with a stable knockout of the enhancer via CRISPR-Cas9-mediated genome editing. We targeted an approximately 400bp region harboring rs6024870 with two guide RNAs (gRNAs) flanking both ends of the putative enhancer. We then utilized a lentiviral vector to introduce the gRNAs, as well as the tracrRNA scaffold, Cas9 gene, and mCherry as a positive selection marker into WT HMC3 cells. mCherry+ cells were sorted by fluorescence-activated cell sorting, and single cell clones were obtained and assessed for deletion by genotyping PCR and Sanger sequencing. We then isolated three individual enhancer-deleted clones that were validated for the desired enhancer deletion, as well as three mock clones transduced with lentivirus containing the tracrRNA scaffold and Cas9 gene, but no targeting gRNA sequence (**Figure 2C**).

To identify transcriptional changes resulting from the rs6024870 enhancer-knockout (KO), we performed bulk RNA sequencing of two mock and three homozygous enhancer-KO clones. Three technical replicate libraries were prepared for each clone and sequenced on the Illumina platform. PCA and heatmap clustering showed that technical replicates clustered together with no batch effects (**Suppl. Fig. 2**). We therefore merged the replicates and performed differential gene expression analysis between the enhancer-KO and mock conditions using FDR < 0.05 and abs(logFC) < 0.58 cutoffs, which yielded 75 differentially expressed genes (47 upregulated and 28 downregulated; **Figure 3A and Suppl. Table 5**). While *RTFDC1* did not reach the threshold for significance, it was downregulated by a fold change of 0.82. We performed Gene Set Enrichment Analysis^52,53^ with GSEA using the Reactome pathways (leveraging all 21,051 genes used in the differential expression analysis ranked by log_2_FC*-log_10_ *P*-value) and identified five upregulated (**Table 2**) and 105 downregulated pathways (**Table 3 and Suppl. Table 6)**. The five upregulated pathways in enhancer-KO cells were related to inflammation (“interferon alpha and beta signaling”, “interleukin 10 signaling”, and “purinerginc signaling in Leishmaniasis infection”) and cell death (“regulated necrosis” and “TP53 regulated transcription of cell death genes”), while the downregulated pathways varied, including “mRNA splicing”, “unfolded protein response”, and “double strand break repair” among the top dysregulated pathways.

**Figure 3:**
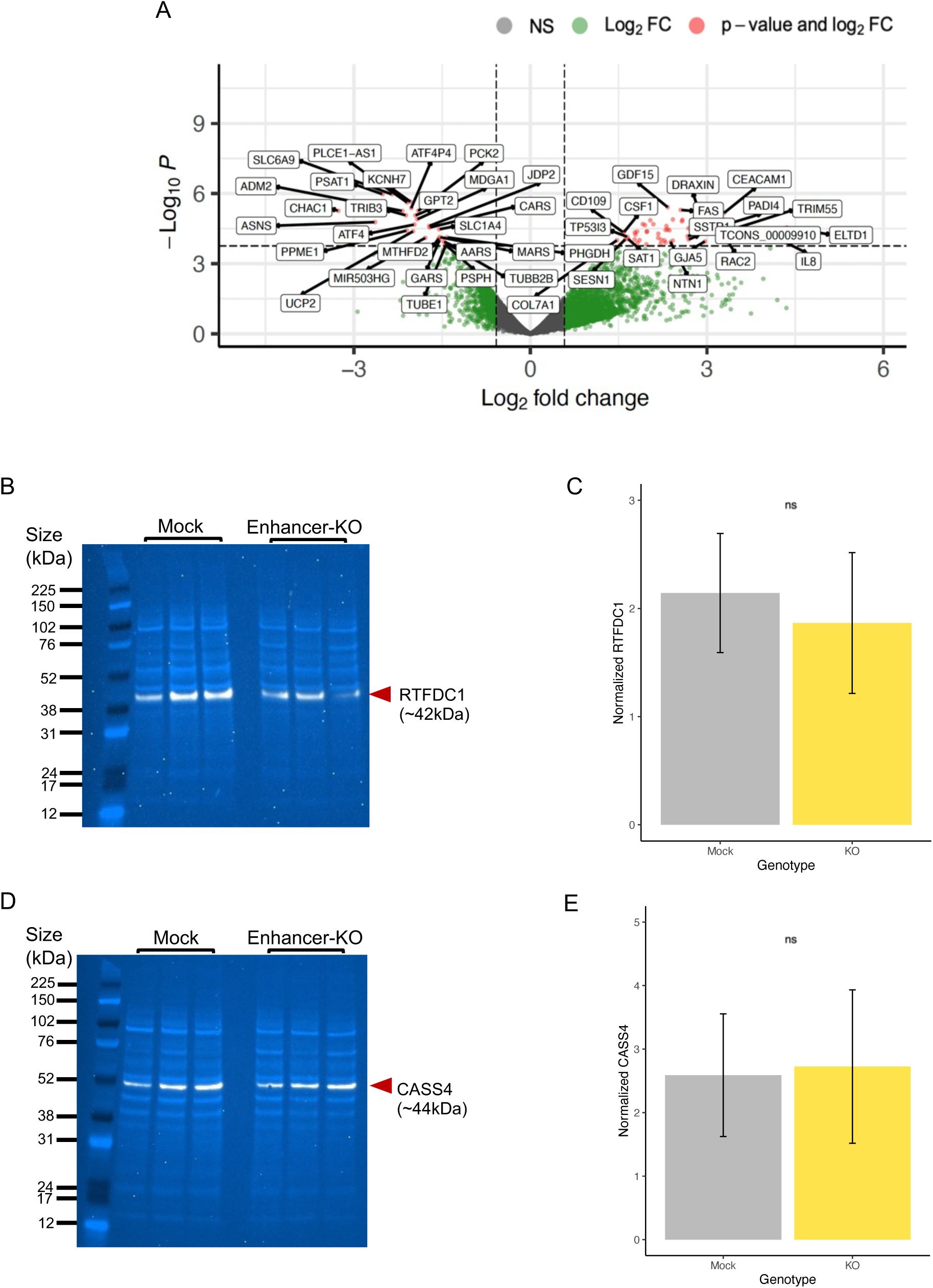
Molecular characterization of rs6024870-enhancer KO HMC3 clones. **(A)** Volcano plot showing differentially expressed genes between 3 homozygous KO and 2 control mock clones. **(B)** Immunoblot for RTFDC1 in mock and enhancer-KO clones. White band: RTFDC1. Light blue bands: whole protein stain. **(C)** Quantification of RTFDC1 immunoblots in mock and enhancer-KO clones. For each genotype, N=6 (3 clones, 2 protein samples per clone). Bars represent sample mean, error bars represet +/-1 standard deviation of the mean. *P*=0.44 by Student’s 2-sample T-test. **(D)** Same as B, but for CASS4. **(E)** Same as C, but for CASS4. *P*=0.83 by Student’s 2-sample T-test.

**Table 2:** Top up-regulated pathways assessed by GSEA.

| GENE SET NAME | GENE SET<br>SIZE | NORMALIZED<br>ENRICHMENT SCORE | FDR q-val |
| --- | --- | --- | --- |
| INTERFERON_ALPHA_BETA_SIGNALING | 64 | 1.8753045 | <0.0001 |
| INTERLEUKIN_10_SIGNALING | 34 | 1.756662 | 0.01154514 |
| PURINERGIC_SIGNALING_IN_LEISHMANIASIS_INFECTION | 26 | 1.7611668 | 0.01500235 |
| TP53_REGULATES_TRANSCRIPTION_OF_CELL_DEATH_GENES | 42 | 1.7284315 | 0.02421465 |
| REGULATED_NECROSIS | 53 | 1.7144489 | 0.02906841 |

**Table 3:**
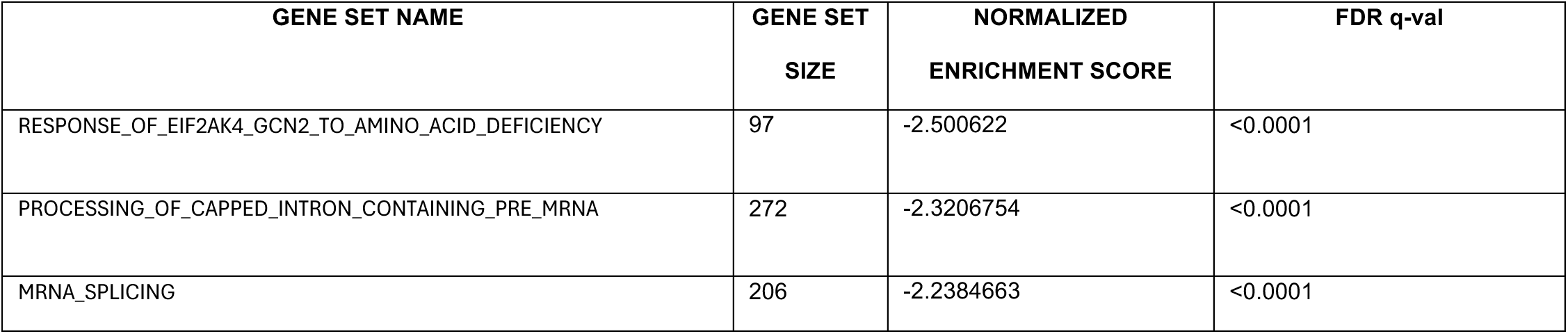
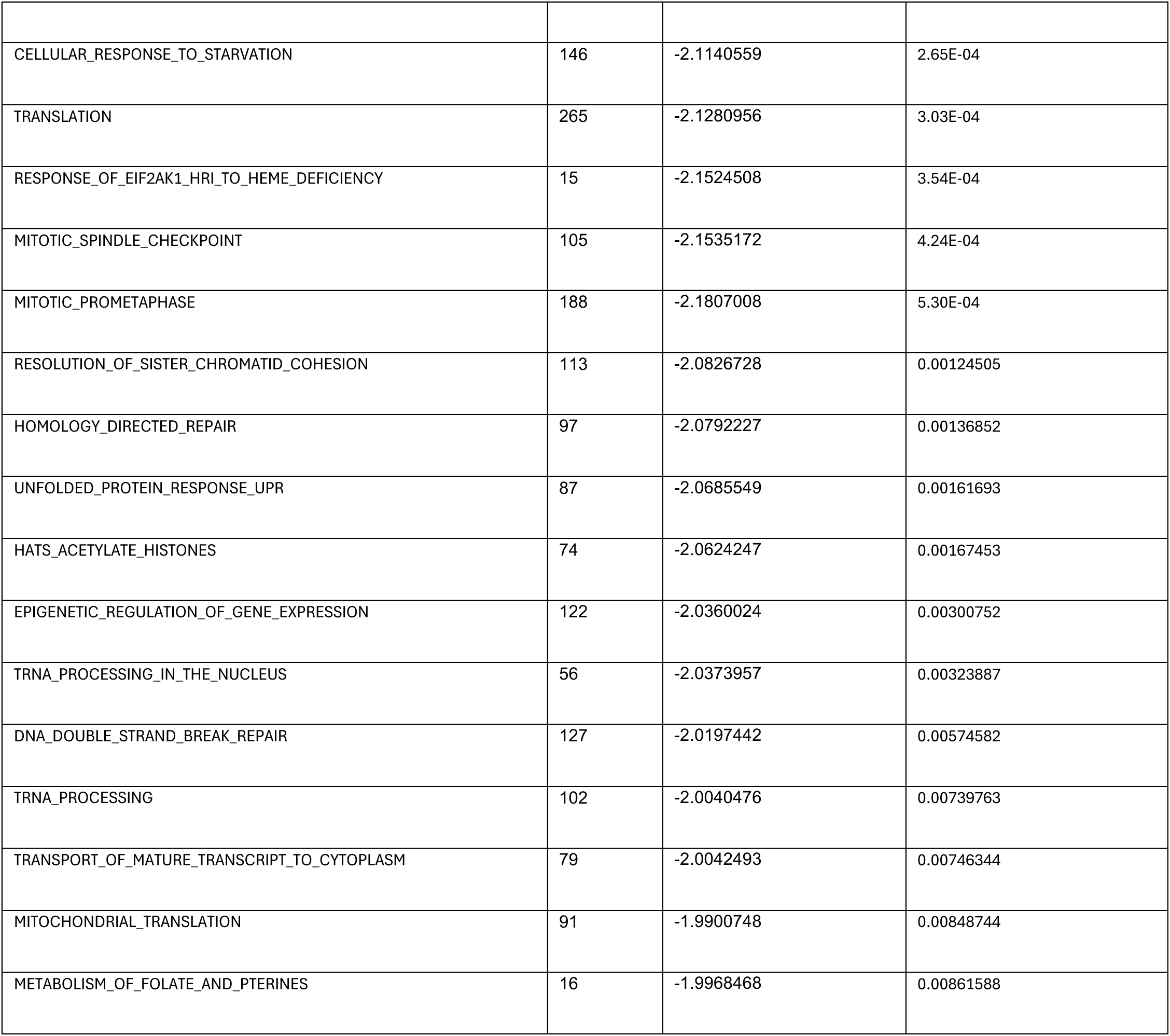
Top down-regulated pathways assessed by GSEA.

To determine if downregulation of *RTFDC1* was detectable at the protein level, we performed immunoblotting for RTFDC1 in enhancer-KO and mock clones. Enhancer-KO clones showed a directional trend of reduced RTFDC1 expression, with an average fold change of 0.72 (normalized to whole protein stain) compared to mock clones, albeit not achieving statistical significance (**Figures 3B and C, Suppl. Figure 3A**), consistent with the direction-of-effect and size-of-effect identified by RNA-seq. We also blotted for CASS4 in the same samples; CASS4 expression was slightly upregulated with an average fold change of 1.11, again albeit not achieving statistical significance (**Figures 3D and E, Suppl. Figure 3B**). These results suggest a potentially complex regulatory role where the enhancer harboring rs6024870 may influence the expression of *RTFDC1* as well as neighboring genes.

### Deletion of the rs6024870-harboring regulatory element primes HMC3 cells to a pro-inflammatory state

Based on the transcriptional analysis of rs6024870 enhancer-KO cells, we sought to investigate the phenotypic consequences of the top up- and down-regulated pathways on microglial function. Given that three out of five upregulated pathways in rs6024870 enhancer-KO cells compared to mock controls were immune-related, we tested if the enhancer-KO cells secreted elevated basal levels of pro-inflammatory cytokines. Many LOAD patients have an pro-inflammatory cytokine signature present in both the peripheral blood and cerebrospinal fluid that can include interleukin (IL)-6^54^, IL-8^55,56^, IL-1β, and TNF-α^57^. We elected to assess the enhancer-deleted cells for their secretion of both IL-6 and IL-8, as HMC3 cells robustly express these cytokines at the mRNA level, as well as secrete them into their culture media^34,58,59^. ELISA for secreted IL-6 and IL-8 showed significantly higher levels of both cytokines (*P* < 0.01 for both cytokines, Student’s 2-tailed T-test) in enhancer-KO cells when compared to mock transduced cells (**Figure 4A and B**).

**Figure 4:**
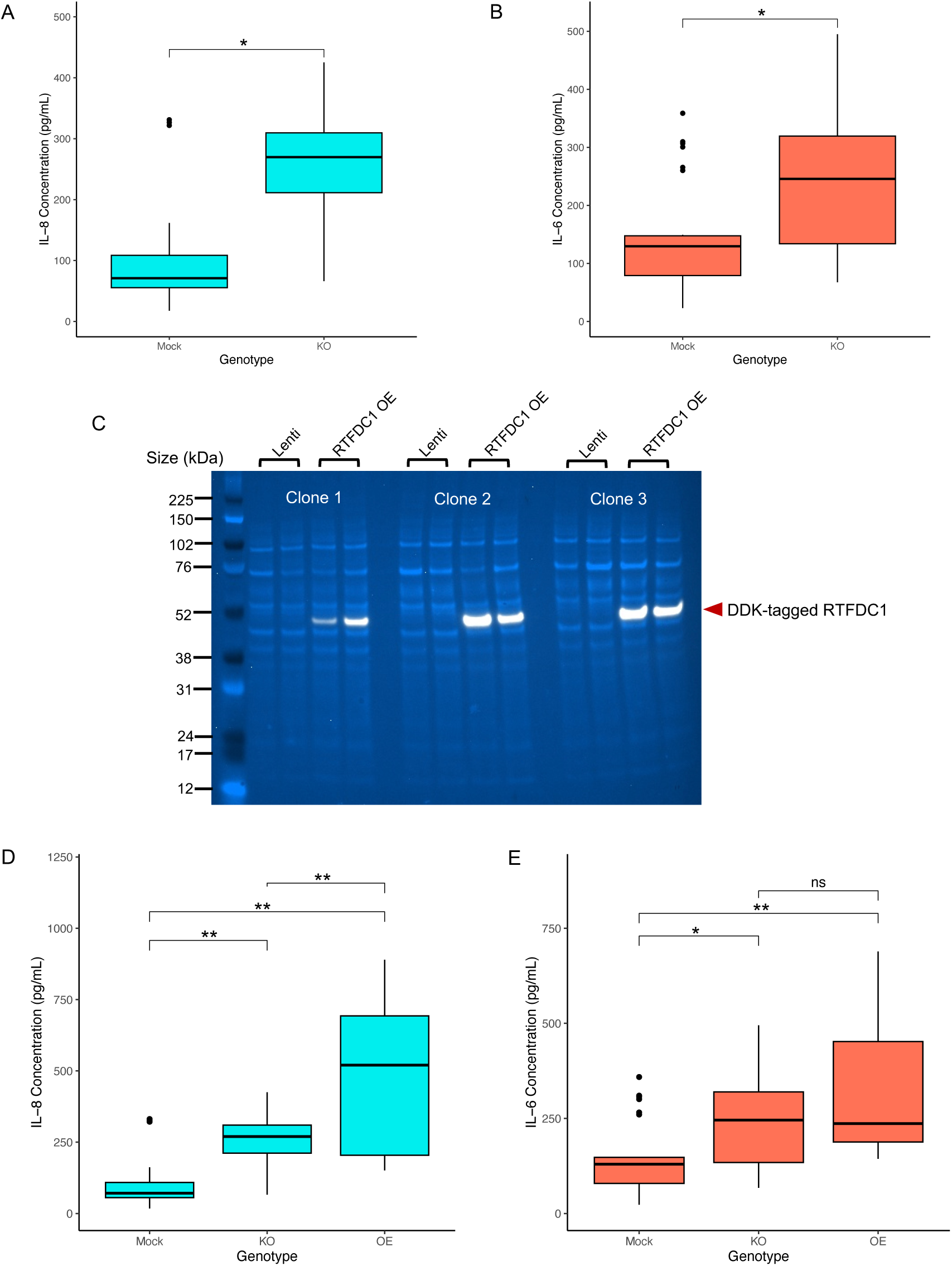
rs6024870-enhancer-KO primes microglia to a pro-inflammatory state. **(A)** Quantification of secreted IL-8 in mock and enhancer-KO cell media. N=27 for each genotype (9 biological replicates per clone, 3 clones per genotype). Boxes represent interquartile range (IQR) with the central bar representing the median (50^th^ percentile), whiskers represent data 1.5 IQR above the 75^th^ percentile or below the 25^th^ percentile, dots represent outliers. * *P*<0.01. **(B)** Same as A, but for secreted IL-6. **(C)** Immunoblot for DDK-tagged lentivirally overexpressed RTFDC1 in all 3 enhancer-KO clones. Lenti: mock transduced. RTFDC1 OE: transduced with DDK-tagged *RTFDC1* ORF. White band: DDK-tagged RTFDC1. Light blue bands: whole protein stain. **(D)** Quantification of secreted IL-8 in mock, enhancer-KO, and RTFDC1 overexpression enhancer-KO clones. N=18 for RTFDC1 overexpression genotype (6 biological replicates per clone, 3 clones per genotype). ** *P*<0.001. **(E)** Same as **D**, but for secreted IL-6. * *P*<0.01, ** *P*<0.001.

To determine if the pro-inflammatory phenotype observed in the enhancer-KO cells could be rescued by reintroducing *RTFDC1*, we utilized a lentiviral overexpression system of *RTFDC1*. We transduced enhancer-KO cells with a lentiviral overexpression vector containing a DDK-tagged *RTFDC1* open reading frame, and selected successfully transduced cells by 7 days of puromycin selection. We then validated the overexpression efficiency in each individual enhancer-KO clone by immunoblot for DDK and RTFDC1 **(Figure 4C and Suppl. Figure 4 and Suppl, Figure 5)**. Unexpectedly, *RTFDC1* overexpression enhancer-KO cells secreted significantly more IL-8 than their enhancer-KO counterparts (*P*<0.001, one-way ANOVA with Tukey’s HSD test, **Figure 4D**), while the same cells did not significantly differ in secretion of IL-6 *(P*=0.198, **Figure 4E**). We noted that the difference in quantity of both secreted cytokines was not driven by the lentiviral overexpression system itself (**Suppl. Figure 6**). These data suggest that overexpression of *RTFDC1* did not rescue the pro-inflammatory phenotype, and may itself lead to increased IL-8 secretion, and that increased secretion of IL-6 in enhancer-deleted clones is independent of *RTFDC1* expression. These results, together, indicate that the rs6024870-harboring enhancer is necessary to suppress secretion of cytokines in the HMC3 cell line, but does not solely function through RTFDC1 in this process.

### Deletion of the rs6024870-harboring enhancer dysregulates the DNA damage response

As ‘DNA double strand break repair’ was one of the top downregulated pathways in the rs6024870-enhancer KO cells, we sought to investigate defects of DNA damage response in these cells. To assess this, we employed an immunofluorescence-based assay for accumulation of the universal double-strand break (DSB) marker phosphorylated serine-139 on histone H2AX (γH2AX), an epigenetic mark associated with accrual of DSBs and recruitment of DNA damage response proteins to genotoxic damage^60^ (**Figure 5A, Suppl. Figure 8A**). We induced DSBs with etoposide, washed out the etoposide, and then stained fixed cells for γH2AX accumulation immediately (hour 0) and 24 hours after DSB-induction to determine if enhancer-deleted cells exhibited a defect in DSB resolution. Enhancer-KO cells showed little difference in γH2AX staining from mock controls at either baseline (control) or hour 0 post-DSB induction. However, enhancer-KO cells had significantly higher levels of γH2AX fluorescence, and thus less resolution of DSBs, at the 24 hour time point than did the mock controls (*P*<2.2×10^−16^, two-way ANOVA (genotype by condition) with Tukey’s HSD, **Figure 5B**). Quantification of γH2AX clearance from hour 0 to hour 24, normalized to control fluorescence levels, showed that mock transduced cells cleared 41.2% of γH2AX fluorescence, whereas enhancer-KO cells only cleared 20.5%.

**Figure 5:**
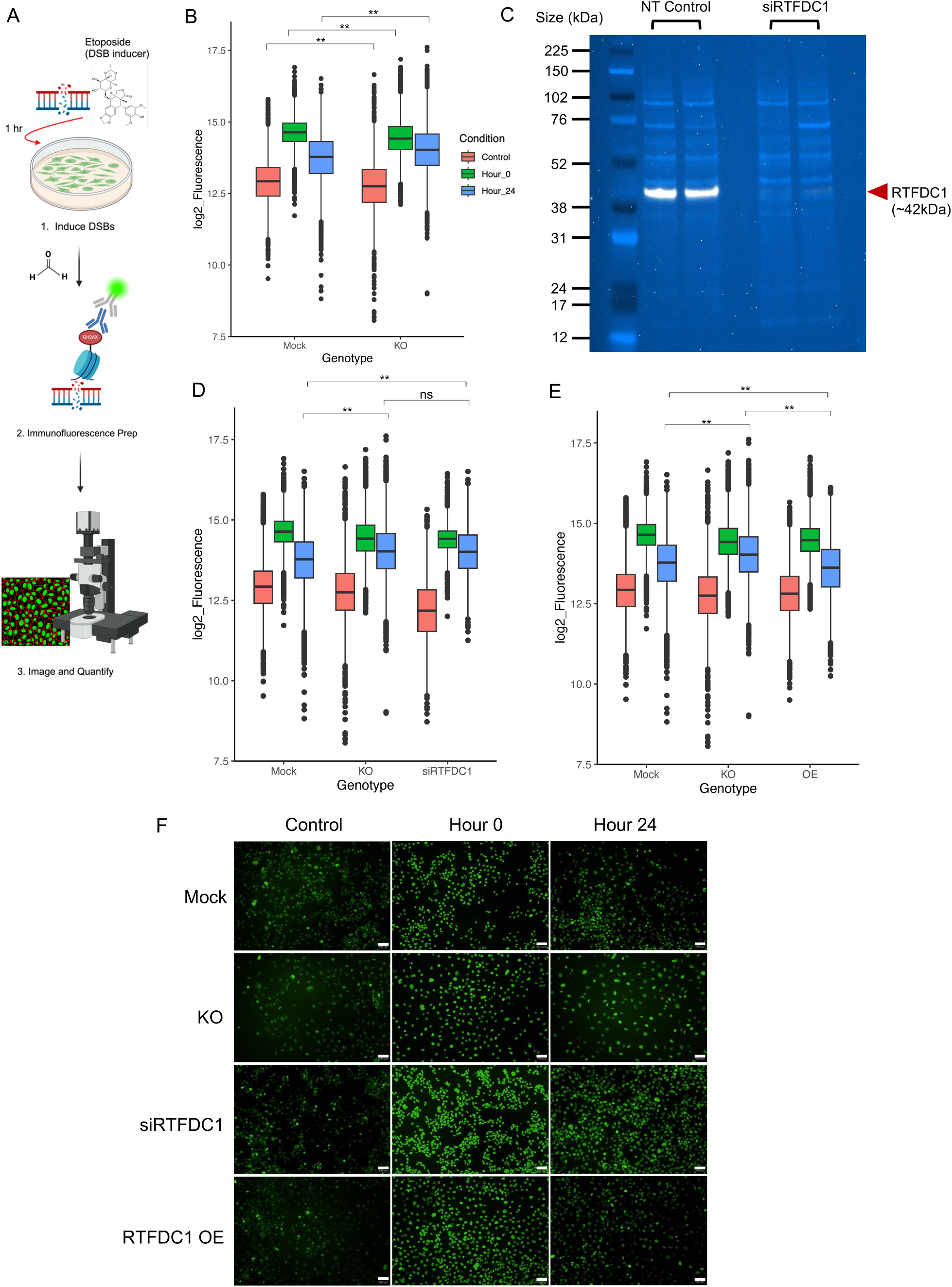
Reduction of *RTFDC1* expression dysregulates the DNA damage repair response. **(A)** Visual schematic of γH2AX staining experimental design. **(B)** Quantification of γH2AX fluorescence in mock and enhancer-KO cells across 3 time periods: no etoposide control (red), 0 hours after etoposide removal (green), and 24 hours after etoposide removal (blue). Boxes represent interquartile range (IQR) with the central bar representing the median (50^th^ percentile), whiskers represent data 1.5 IQR above the 75^th^ percentile or below the 25^th^ percentile, dots represent outliers. For each genotype, N=12 wells (4 wells per clone, 3 clones per genotype). ** *P*<2.2×10^−16^ by two-way ANOVA (genotype by condition) with Tukey’s HSD. **(C)** Immunoblot for RTFDC1 in WT HMC3 cells transduced with either non-targeting control siRNA (NT Control) or *RTFDC1* siRNA pool (siRTFDC1). Each lane represents an individual transfection. White band: RTFDC1. Light blue bands: whole protein stain. **(D)** Quantification of γH2AX fluorescence in mock, enhancer-KO, and siRNA RTFDC1-depleted cells (siRTFDC1) across same time periods as **B**. For siRNA RTFDC1-depleted cells, N=4 wells (2 wells per siRNA transfection, 2 transfections total). ** *P*<2.2×10^−16^ by two-way ANOVA (genotype by condition) with Tukey’s HSD. **(E)** Quantification of γH2AX fluorescence in mock, enhancer-KO, and RTFDC1-overexpression cells (OE) across same time periods as **B**. For overexpression cells, N=12 wells (4 wells per clone, 3 clones per genotype). ** *P*<2.2×10^−16^ by two-way ANOVA with Tukey’s HSD. **(F)** Representative images of γH2AX fluorescence (green) across all conditions and timepoints. All images 10X magnification. White bars represents 100μm.

To investigate whether this phenotype was dependent on *RTFDC1* levels, we employed *RTFDC1* siRNA knockdown. We transfected WT HMC3 cells with a pool of siRNAs targeting RTFDC1, and evaluated knockdown efficiency 72 hours post-transduction via immunoblot (**Figure 5C, Suppl. Figure 7**). *RTFDC1* knockdown cells exhibited a 95% decrease in RTFDC1 protein expression. Indeed, these *RTFDC1* knockdown cells also yielded significantly elevated levels of γH2AX at the 24 hour time point when compared to mock transduced controls (*P*<2.2×10^−16^, two-way ANOVA (genotype by condition) with Tukey’s HSD), and did not differ significantly from enhancer-deletion cells, resolving 18.5% of γH2AX fluorescence (**Figure 5D**). We then assessed whether overexpression of *RTFDC1* in enhancer-KO cells was sufficient to rescue the reduction of γH2AX fluorescence clearance using our lentiviral *RTFDC1* overexpression system (**Figure 4C**). Indeed, over-expression of *RTFDC1* increased the clearance of γH2AX fluorescence from hour 0 to hour 24 to 41.6%, and the mean fluorescence at hour 24 was in fact lower than even mock transduced controls (*P*<2.2X10^−16^, two-way ANOVA (genotype by condition) with Tukey’s HSD, **Figure 5E**). The γH2AX fluorescence differences in both siRNA RTFDC1-depleted and lentiviral RTFDC1-overexpression cells was not driven by either the siRNA transfection or lentiviral overexpression systems (**Suppl. Figure 8B and C**). Representative images of fluorescence across all conditions and timepoints are displayed in **Figure 5F**. This data suggests that the dysregulation in DNA damage response in the enhancer-KO HMC3 cells is mediated, at least in part, by *RTFDC1*.

### Cumulative evidence of rs6024860 influences *RTFDC1* expression in microglial cells

Next, we sought to validate the enhancer activity of the region harboring rs6024870 in the HMC3 human microglial cell model and investigate whether the enhancer’s activity was modulated by rs6024870 specifically. To assess this, we used a dual-luciferase assay in the HMC3 human microglia-like cell line. We cloned the approximately 1.5kb endogenous human promoter of *RTFDC1* upstream of the *Photinus pyralis luc2* gene. Immediately upstream of the *RTFDC1* promoter, we cloned in a 1,031bp region containing either rs6024870 allele, which displayed enhancer marks of both open chromatin and H3K27ac. Utilizing site-directed mutagenesis, we generated plasmids that differed at only the major (G) and minor (A) allele of rs6024870 (**Figure 6A**). These plasmids, along with negative and internal controls plasmids, were transfected into WT HMC3 cells, and luminescence of whole-cell lysates was obtained approximately 48 hours later.

**Figure 6:**
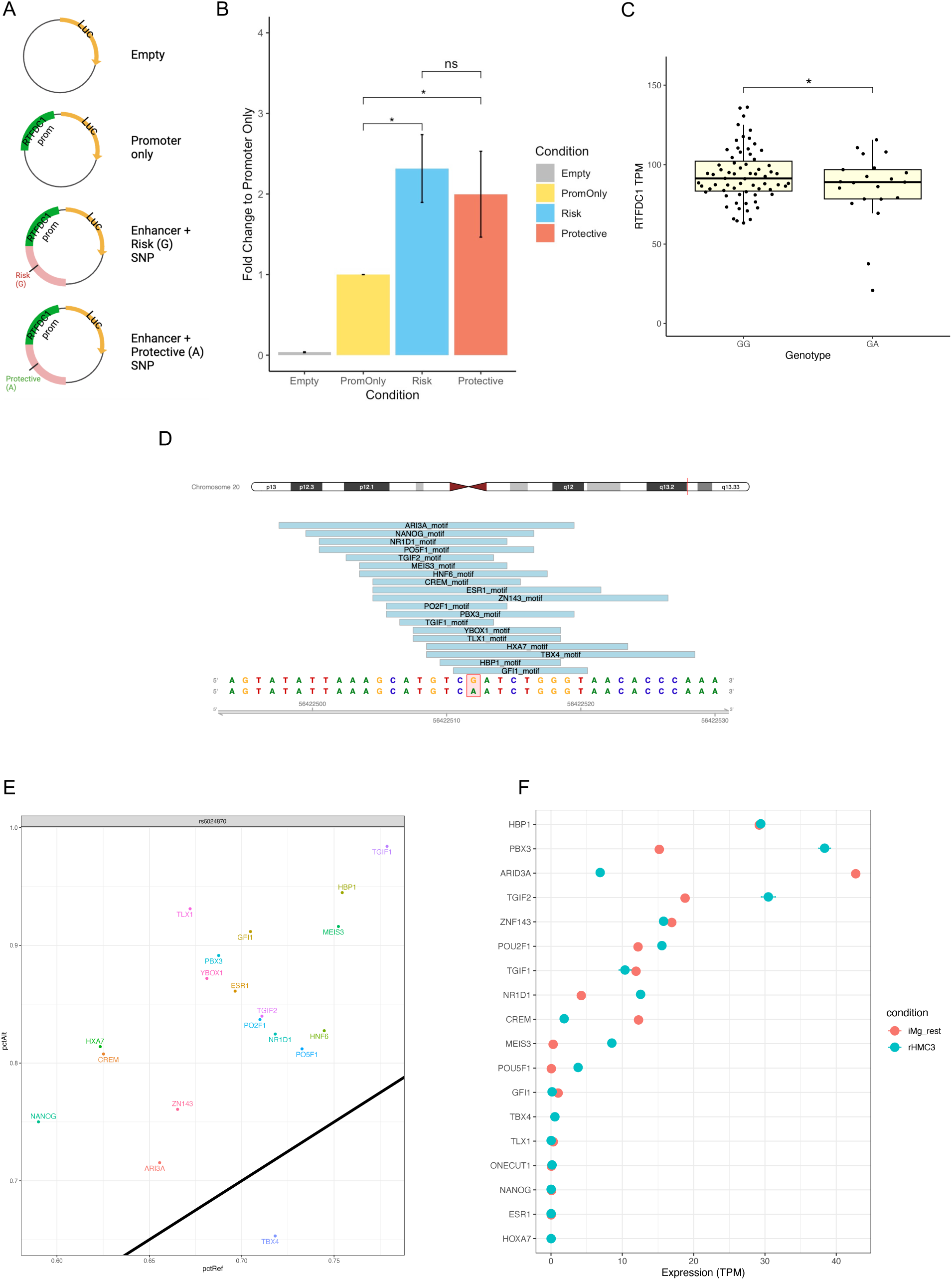
rs6024870 influences *RTFDC1* expression in microglial cells. (A) Schematic of plasmid design used for luciferase experiments. (B) Luciferase assay luminescence fold change of rs6024870-harboring regulatory element in HMC3 cells. N=7 biological replicates per condition. (C) Expression analysis of *RTFDC1* TPMs from freshly harvested microglial cells from the FreshMicro study^40^. * *P*=0.0385 by Student’s 1-tailed T-test. (D) Transcription factor motifs overlapping rs6024870. (E) Transcription factor motif affinity analysis for rs6024870 genotype with MotifBreakR. Black diagonal line represents no change in predicted binding affinity in the presence of the protective (A) allele. Any motifs above the diagonal are predicted to have increased binding affinity in the presence of the protective allele, while motifs below the diagonal are predicted to have weakened binding affinity in the presence of the protective allele. (F) Expression of transcription factors in iMg (red) and HMC3 (blue).

Luciferase experiments showed that the putative enhancer region harboring the major (G) risk allele increased luciferase expression by approximately 2.3-fold when normalized to *RTFDC1-*promoter-only controls (*P*<0.001, one-way ANOVA with Tukey’s HSD, N = 7 assays). While the enhancer region containing the minor (A) protective allele did show a nominal decrease in luciferase expression, with a 2.0-fold expression change compared to promoter-only controls, it did not reach the threshold of significance (**Figure 6B**). These results revealed that the region harboring rs6024870 acts as an enhancer on *RTFDC1* expression in HMC3 cells, and that the minor protective A allele trends towards decreased activity.

Seeking further support of an allele-specific difference on enhancer activity at rs6024870, with the minor allele reducing enhancer activity on *RTFDC1*, we utilized several orthogonal methods. We first leveraged GTEx data to investigate if rs6024870 acted as an eQTL on *RTFDC1*. While we did not find evidence of rs6024870 acting as an eQTL in bulk brain-related tissue, we did observe that rs6024870 acts as an eQTL on *RTFDC1* in thyroid tissue, a tissue rich in macrophages, with a normalized effect size of -0.12 relative to the minor allele (beta *P*-value = 7.67×10^−5^)^41^, directionally consistent with our other findings that the minor protective A allele reduces *RTFDC1* expression.

We then leveraged publicly available patient RNA-seq data of microglia isolated from fresh human brain tissue^58^ to determine whether rs6024870 acted as an eQTL on *RTFDC1* mRNA transcript abundance. We identified from the dataset 70 individuals homozygous for the major allele (GG), 21 heterozygous individuals (GA), and only 2 individuals homozygous for the minor allele (AA). This is consistent with expected Hardy Weinberg genotype proportions, as the minor A allele frequency (MAF) of rs6024870 is 0.058 in European ancestry. We therefore elected to focus our analysis only on GG and GA individuals. Given our *a priori* hypothesis that the minor allele A reduces *RTFDC1* expression, we performed a Student’s 1-tailed T-test on the expression data for GG and GA individuals, removing all TPM outliers +/-3 standard deviations of the mean, and found that GG individuals had on average 93.40 TPM of *RTFDC1*, significantly higher than GA individuals, which had on average 85.30 TPM (*P*=0.0385, Student’s 1-tailed T-test, **Figure 6C**).

Many enhancers operate via binding with transcription factors (TFs) to facilitate contact with promoters^61^. To identify if TF binding can be affected by the presence of the rs6024870 risk allele, we performed TF motif analysis on the open chromatin region harboring rs6024870. We identified 19 different TF motifs overlaying rs6024870 (**Figure 6D, Suppl. Figure 9**). Binding affinity analysis with motifBreakR showed that 18 of 19 of these motifs were strengthened in the presence of the minor allele (**Figure 6E**). The majority of these TFs are expressed in our microglial models (**Figure 6F**), and are reported to act as transcriptional repressors, such as TGIF2^62^, HBP1^63^, and NR1D1^64^, which is consistent with the observed trend of decreased expression of *RTFDC1* in the presence of the minor protective A allele. While further work is necessary to validate these observations, these suggest that rs6024870 may function to restrain enhancer activity.

Collectively, we observed cumulative evidence of consistent directionality that the protective allele at rs6024870 reduces expression of *RTFDC1* in microglial cells.

## DISCUSSION

In this study, we utilized a genomics-based variant-to-gene mapping approach in two different model systems of human microglia, iPSC-derived microglia (iMg) and the HMC3 cell line, to map physical contacts between LOAD GWAS-implicated variants and their associated regulatory elements with gene promoters.

Recently, two studies using proximity-ligation assays in either human primary or iPSC-derived microglia have reported 3D genomic interactions between GWAS-associated variants and gene promoters in LOAD^22,65^. However most Hi-C-based techniques are at too low a resolution to detect specific GWAS variant-promoter contacts due to their utilization of a relatively infrequent 6-cutter enzyme (HindIII)^66^. Additionally, ChIP-based proximity ligation assays, such as PLAC-seq^67^, rely on the use of antibodies to pull down pre-specified regions associated with certain histone marks that can demarcate putative enhancer (H3K27ac) or promoter (H3K4me3) regions. Our study improves on prior studies by utilizing a relatively frequent 4-cutter restriction enzyme, DpnII, which generates smaller fragments with higher resolution across the entire genome^46^. Additionally, our Capture-C library assessed a comprehensive set of primary and alternative human gene promoters for both coding and non-coding genes, allowing us to probe all known promoters’ interactions without any prior knowledge of histone acetylation status. Furthermore, it has been recently shown that genes near GWAS signals are enriched for key functional annotations and are under strong selective constraint, while genes near eQTLs lack most functional annotations and show relaxed constraint^68^; importantly, and in contrast to eQTL-driven methods, our 3D genomic-informed approach is not constrained by these detection differences. This in turn enabled us to generate unbiased, hypothesis-free interaction maps of LOAD GWAS-associated variants with putative effector gene promoters in our microglial cell model systems.

Our combination of promoter-focused Capture-C and ATAC-seq data identified 33 LOAD GWAS-associated variants contacting the promoters of 32 putative effector genes, 24 of which code for a protein, with 5 targets overlapping in both cellular models (*AURKA*, *CSTF1*, *MIR4487*, *RTFDC1*, and *TCONS_00028450*). From this list we observed consistent evidence of a candidate regulatory genomic region exhibiting enhancer epigenetic signatures specifically in microglial cells at the *CASS4* GWAS locus, harboring the LOAD-associated variant, rs6024870 (r^2^ = 0.97 with its sentinel variant, rs6069737). This region contacted the promoter of *RTFDC1* in both cellular settings and its role in LOAD susceptibility remained to be defined. What is known to date is that in *S. Pombe* the removal of *RTFDC1* (also known as *RTF2*) from stalled replication forks during S phase DNA replication is essential to maintain replication stress and genome integrity^69^. However, more recent work has also shown that in humans, RTFDC1 is required for appropriate localization of RNAseH2 to the replication fork to remove ribonucleotides from nascent DNA via the ribonucleotide excision repair pathway^70^.

Our characterization of the genomic region harboring rs6024870 in HMC3 cells indicated that this region acts as a microglial specific enhancer, and our CRISPR-Cas9 induced ∼400bp deletion of this enhancer provided variant-to-function evidence for its role in regulating both microglial inflammation (upregulation of inflammatory pathways, increased secretion of IL-8 and IL-6) and DNA damage response (downregulation of DNA repair pathways, loss in ability to resolve DNA DSBs as assayed by fluorescent γH2AX accumulation).

While the exact effector genes at this locus remain to be elucidated, our study suggests a role for *RTFDC1*, as indicated by our observation that rs6024870 acts as a modest eQTL on *RTFDC1* expression in microglia from human brains, with the minor A allele (protective against LOAD) associated with reduced expression levels. This direction of effect is consistent with a recent proteome-wide association study in LOAD showing that the brains of LOAD patients from the ROS-MAP cohort presented with increased levels of RTFDC1 when compared to unaffected controls^71^. This is also consistent with *RTFDC1* overexpression in enhancer-KO cells exhibiting increased secretion of cytokine IL-8, which is a pro-inflammatory phenotype thought to be detrimental in LOAD. In contrast, secrection of IL-6 was not affected by RTFDC1 overexpression, suggesting that this phenotype is independent of *RTFDC1*.

In HMC3 cells, CRISPR-Cas9-mediated KO of the rs6024870-harboring enhancer region resulted in a DNA double strand break defect. While we could recapitulate this phenotype by knocking down *RTFDC1* expression in WT cells via siRNA and by lentivirally overexpressing *RTFDC1* in the KO cells, which strongly suggest that this effect is, at least in part, mediated by RTFDC1, we were unable to detect a significant difference in RTFDC1 levels in our enhancer-KO HMC3 cells, both at the RNA or the protein level. This may be due to technical difficulties such as insufficient sensitivity of our assays to detect a relatively subtle change in its expression levels, but could also suggest that other effector genes at this locus besides *RTFDC1* are playing a role in this phenotype. While our variant-to-gene mapping datasets implicated a physical contact between the regions harboring rs6024870 and the promoter of *RTFDC1*, the activity-by-contact model predicts that this SNP also contacts multiple other genes within the same TAD, including *CASS4, CSTF1*, *AURKA, FAM210B,* and *TFAP2C* in THP1 cells (an immortalized human monocyte cell line derived from an acute monocytic leukemia patient)^72,73^. This is consistent with the observation that a single enhancer may contact and regulate several promoters, a phenomenon previously seen in macrophages^74^. The enhancer identified in our study likely regulates multiple genes, including *RTFDC1*, either simultaneously or at different timepoints throughout the lifespan of the microglia, where the regulation of these additional genes may contribute to the pro-inflammatory and the DNA-repair phenotypes we observe in our enhancer-deleted cells.

Future efforts are warranted to characterize the full regulatory profile of this enhancer in our microglial model systems in order to better understand both how this enhancer functions in relation to LOAD pathology and the allele-specific effects of rs6024870 on its activity. LOAD is a disease of aging, and the protective allele at rs6024870 may confer its protective benefits only later in life. The relatively low MAF of rs6024870 suggests that the rs6024870 protective allele is deleterious earlier in life, due to the evolutionary recency of expansion of life expectancy. In fact, the strongest protective allele against LOAD, *APOEρ.2*, has the lowest frequency of the three alleles at the *APOE* locus^75^. In addition to conferring a protective advantage against LOAD and other dementias, individuals homozygous for the *APOEρ.2* allele also have an increased risk of reduced fertility^76^, as well as poorer cancer outcomes^77^; therefore, one could hypothesize that is the reason for its low frequency in the population, as its benefits do not manifest until much later in life. Taken together, one hypothesis is that the minor allele of rs6024870 confers a protective advantage to older individuals, with its low allele frequency potentially driven by more deleterious effects in younger individuals.

To the best of our knowledge, we are the first to perform physical variant-to-gene mapping in the HMC3 cell line. Our characterization of the HMC3 3D genomic architecture, chromatin accessibility, and transcriptional profile will help to further establish it as a useful model system to study microglial function in LOAD, as well as other neurological diseases. However, we acknowledge the limitations of working with an immortalized cell line. HMC3 cells were originally immortalized by transfecting human embryonic microglial cells with a plasmid encoding the cDNA of the viral SV40 large T antigen^78^. While these cells maintain much of the characteristic antigenic profile of primary human microglia, including IBA1 at rest and CD68 in activated conditions, they do not secrete LOAD-relevant tumor necrosis factor α (TNF-α) or IL1-β^78^, which limited the cytokines and chemokines we could assay in our study. Future work is warranted to further validate our enhancer-deletion in the iMg model system to characterize primary microglia characteristics, with the ultimate goal of generating iMg lines harboring a single base pair change at rs6024870 to characterize the allele-specific effect of this variant in a model closer to the *in vivo* state of human microglia.

Overall, we have physically mapped LOAD-associated GWAS-implicated variants to their putative regulatory elements and corresponding candidate effector genes in two human microglial systems, investigated the function of the rs6024870-harboring microglial enhancer in HMC3 cells using a CRISPR-Cas9 mediated precise deletion, and provided cumulative evidence for a role of rs6024870 and *RTFDC1* in LOAD pathogenesis, including a novel role for *RTFDC1* in DNA double strand break repair. Through these efforts, we have also generated multiple genomic datasets, including ATAC-seq, RNA-seq, and Promoter-focused Capture-C, for both the HMC3 cell line and iMg cells, which will serve as a resource for further study of microglial genetic involvement in LOAD and other neurological diseases. With our variant-to-function approach we implicate both inflammation and DNA damage repair as relevant microglial-specific pathways to the pathology of LOAD, which will help to focus future studies in identifying genes that can be effectively targeted for the development of therapeutic interventions that can in turn improve LOAD patient outcomes.

## METHODS

All primers and small RNAs utilized in this study are detailed fully in **Supplementary Table 7**.

### Loci Analyzed

We extracted all sentinel SNPs for the significant loci reported in the recent GWAS meta analyses from Bellenguez *et al.* and Wightman *et al*^4,31^. for a total of 111 SNPs (83 from Bellenguez; 38 from Wightman; 10 index SNPs were in common). We retained one representative SNP from the 10 pairs that were in high LD (r^2^>0.7), resulting in 101 SNPs under consideration in total. We obtained all proxy SNPs in high LD (r^2^>0.7) with these sentinel SNPs using SNiPA^32^, selecting the European panel of the 1000 Genomes phase 3 v.5 database.

### Cell Culture

HMC3 (CRL-3304) and 293T (CRL-3216) cells were purchased from ATCC. HMC3 cells were grown in Minimal Essential Media (Gibco) supplemented with 10% heat-inactivated FBS (Gibco) and 1X antibiotic-antimycotic (Gibco). 293T cells were grown in Dulbecco’s Minimal Essential Media (Gibco) supplemented with 10% heat-inactivated FBS and 1X antibiotic-antimycotic. Cells were maintained at 37.0°C using standard cell culture conditions. iPSCs obtained from the CHOP Stem Cell Core (CHOP WT10 line^79^) were differentiated into common myeloid progenitors (CMPs) according to published protocol^80^. CMPs were then differentiated to iMicroglia according to published protocol^33^.

### Mycoplasma Contamination Testing

1mL of Normal growth media was removed from individual cell culture dishes and centrifuged at 15,000rpm for 5 minutes at room temperature. Supernatants were tested for the presence of mycoplasma using the InvivoGen MycoStrip^TM^ Detection kit according to manufacturer’s protocol with no alterations. The kit included mycoplasma DNA as a positive control, and sterile PBS was used as a negative control.

### Cell Fixation for Chromatin Capture

The protocol used for cell fixation was similar to previous methods^20^. Cells were fixed in culture plates for 10 minutes at room temperature on a rocking platform using 270µL of formaldehyde per 10mL growth media. The reaction was quenched by adding 1.5mL 1M cold glycine (4 °C). Volumes were adjusted based on culture vessel size and media volume for each cell type. Cells were collected from plates using a cell scraper and pooled into a single-cell suspension of 10^7^ cells. Fixed cells were centrifuged at 1000rpm for 5 minutes at 4 °C and the supernatant was removed. The cell pellets were washed in 10mL cold PBS (4 °C) followed by centrifugation as above. Supernatant was removed and cell pellets were resuspended in 5mL of cold lysis buffer (10mM Tris pH8, 10mM NaCl, 0.2% NP-40 (Igepal) supplemented with protease inhibitor cocktails). Resuspended cells were incubated for 20 minutes on ice, centrifuged as above, and the lysis buffer was removed. Finally, cell pellets were resuspended in 1mL fresh lysis buffer, transferred to 1.5mL Eppendorf tubes and snap frozen in ethanol and dry ice. Cells were stored at −80 °C until they were thawed for 3C library generation.

### 3C Library Generation

For each library, 10^7^ fixed cells were thawed on ice, pelleted by centrifugation and the lysis buffer removed. The cell pellet was resuspended in 1mL dH_2_O supplemented with 5µL protease inhibitor cocktail (200X) and incubated on ice for 10 minutes, followed by centrifugation and removal of supernatant. The pellet was then resuspended in dH_2_O for a total volume of 650µL. Control 1 (50µL) was removed at this point and remaining samples were divided into 6 tubes. NEBuffer DpnII (1X), dH_2_O, and 20% SDS were added and samples were incubated at 1000rpm for 1 hour at 37 °C in a MultiTherm (Sigma-Aldrich). Following the addition of Triton X-100 (concentration, 20%), samples were incubated an additional hour. Next, 10µL of DpnII (50 U/µL) (NEB) was added and incubated for 4 hours at 37 °C. An additional 10µL DpnII was added and digestion was left overnight. The next day, another 10µL of DpnII was added and incubated for a further 3 hours. 100µL of each digestion reaction was removed to generate control 2. Finally, samples were incubated at 1000rpm for 20 minutes at 65 °C to inactivate the DpnII and placed on ice for 20 additional minutes.

Next, the digested samples were ligated with 8µL T4 DNA Ligase (HC ThermoFisher, 30 U/µL) and 1X ligase buffer at 1000rpm overnight at 16 °C in the MultiTherm. The next day, an additional 2 µL T4 DNA ligase was spiked in to each sample and incubated for 3 more hours. The ligated samples were then de-crosslinked overnight at 65 °C with Proteinase K (Invitrogen) and the following morning incubated for 30 minutes at 37 °C with RNase A (Millipore). Phenol-chloroform extraction was then performed, followed by ethanol precipitation overnight at −20 °C and a 70% ethanol wash. Samples were resuspended in 300µL dH_2_O and stored at −20 °C until Capture-C. Digestion efficiencies of 3C libraries were assessed using control 1 (undigested) and control 2 (digested) by gel electrophoresis on a 0.9% agarose gel and quantitative PCR (SYBR green, Thermo Fisher).

### High-resolution promoter focused Capture-C

The 3C libraries were quantified using a Qubit fluorometer (Life Technologies), and 10μg of each library was sheared in dH_2_O using a Qsonica Q800R to an average DNA fragment size of 350bp. Qsonica settings used were 60% amplitude, 30 seconds on, 30 seconds off, 2 minute intervals, for a total of five intervals at 4 °C. After shearing, DNA was purified using AMPureXP beads (Agencourt). Sample concentration was checked via Qubit fluorometer and DNA size was assessed on a Bioanalyzer 2100 using a 1000 DNA Chip. Agilent SureSelect XT Library Prep Kit (Agilent) was used to repair DNA ends and for adaptor ligation following the standard protocol. Excess adaptors were removed using AMPureXP beads. Size and concentration were checked again before hybridization. 1μg of adaptor ligated library was used as input for the SureSelect XT capture kit using their standard protocol and our custom-designed Capture-C library. We leveraged a custom designed capture library (Agilent) targeting all human coding genes plus non-coding transcripts^20^. The quantity and quality of the captured library were assessed by Qubit fluorometer and Bioanalyser using a high sensitivity DNA Chip. Each SureSelect XT library was paired-end sequenced on an Illumina NovaSeq 6000, generating ∼1.6 billion paired-end reads per sample (51bp read length).

### Analysis of Capture-C Data

We performed quality control of the raw fastq files with FastQC. For the iMicroglia samples, the same libraries were sequenced at both 50 and 100bp read length, and the resulting fastq files were merged before processing. Paired-end reads were pre-processed using the HiCUP pipeline and aligned with bowtie2 to the reference genome (hg19). We called significant interactions at 1-DpnII and 4-DpnII fragment resolutions using CHiCAGO with default parameters except for binsize which was set to 2500. For 4-fragment resolution calls, we used artificial .baitmap and .rmap files grouping 4 consecutive DpnII fragments together and using default parameters (except for removeAdjacent which was set to False). The CHiCAGO function peakEnrichment4Features() was used to assess enrichment of genomic features in promoter interacting regions at both 1-fragment and 4-fragment resolution. The UCSC Genome Browser was used to visualize the detected interactions within the context of other relevant functional genomics annotations.

### ATAC-seq library generation

Fresh HMC3 or iMicroglia cells were harvested via Trypsin followed by a series of DPBS wash steps. 50,000 cells from each sample were pelleted at 550 × g for 5 minutes at 4 °C. The cell pellet was then resuspended in 50μl cold lysis buffer (10mM Tris-HCl, pH 7.4, 10mM NaCl, 3mM MgCl2, 0.1% IGEPAL CA-630) and centrifuged immediately at 550 × g for 10 minutes at 4 °C. The nuclei were resuspended in transposition reaction mix (2x TD Buffer (Illumina Cat #FC-121–1030, Nextera), 2.5µl Tn5 Transposase (Illumina, 20034197 Cat #FC-121–1030, Nextera) and Nuclease Free H_2_O) on ice and then incubated for 45 minutes at 37°C. The transposed DNA was then purified using the MinElute Kit (Qiagen), eluted with 10.5μl elution buffer (EB). The transposed DNA was PCR amplified and indexed using the Illumina Nextera Kit (Illumina) and NEBNext High-Fidelity 2x PCR Master Mix (NEB) for 12 cycles to generate each library. The PCR reaction was subsequently purified using AMPureXP beads (Agencourt) and checked for quantity and quality by Qubit (Invitrogen) and Bioanalyzer (Agilent). Libraries were paired-end sequenced (51bp read length at each end) on the Illumina NovaSeq 6000 platform. Open chromatin regions were called using the ENCODE ATAC-seq pipeline (https://www.encodeproject.org/atac-seq/), selecting the resulting IDR optimal peaks (with all coordinates referring to hg19). We define a genomic region open if it has 1bp overlap with an ATAC-seq peak.

### Luciferase Assays

To amplify the 1031bp putative enhancer region enriched in H3K27ac in microglial cells and the harboring rs6024870 major allele (G), we designed PCR primers at the 5’ and 3’ ends of the sequence containing 40bp overhangs for cloning, and amplified the region using the Platinum HotStart PCR mastermix containing Platinum *Taq* polymerase (ThermoFisher Scientific) off of HMC3 genomic DNA as a PCR template. We purified the resulting fragment from a 2% agarose gel using the NucleoSpin Gel and PCR Purification Kit (Macherey-Nagel). We amplified the 1483bp *RTFDC1* promoter sequence off of a promoter plasmid (Genecopoeia HPRM41362-PF02) using primers designed with 40bp overhangs for cloning.

The promoterless pGL4.10[*luc2*] firefly luciferase promoter was purchased from Promega, and linearized using the *XhoI* restriction enzyme at the multiple cloning site. Linearized fragments were visualized on a 2% agarose gel, then extracted and purified using the NucleoSpin Gel and PCR Purification Kit. The purified *RTFDC1* promoter and major allele enhancer were then inserted at the multiple cloning site, upstream of the *luc2* gene, using the Gibson Assembly HiFi HC 1-Step kit (Codex). At the same time, another vector was generated in the same manner containing just the *RTFDC1* promoter to generate the promoter-only control plasmid (PGL4-*RTFDC1*). Assembled products were transformed into NEB Stable Competent *E. coli* cells by heatshock, plated on LB agar plates supplemented with 100μg/mL ampicillin, and grown overnight at 37°C. Individual colonies were then selected and grown out overnight at 30°C, shaking at 250rpm in 5mL LB broth supplemented with 100μg/mL ampicillin. Plasmid DNA was extracted from *E. coli* using the Qiagen QIAPrep Spin MiniPrep Kit, and resulting plasmids were Sanger sequenced at both the 5’ and 3’ ends of the inserted sequences to confirm successful insertion. The final successful plasmid was sent for Oxford Nanopore sequencing by Plasmidsaurus to confirm fidelity of the entire sequence (named PGL4-RTFDC1-rs6024870Major). The purified final plasmid DNA of PGL4-RTFDC1-rs6024870Major was used as a template for site directed mutagenesis to generate the minor allele (A) plasmid (PGL4-RTFDC1-rs6024870Minor) using the NEB Q5 Site-Directed Mutagenesis kit. The final plasmid was sent for Oxford Nanopore sequencing by Plasmidsaurus to ensure successful mutagenesis of the minor allele and to check for any polymerase errors. Stocks of both PGL4-RTFDC1-rs6024870Major and PGL4-RTFDC1-rs6024870Minor, as well as PGL4-*RTFDC1*, the unmodified pGL4.10[*luc2*], and pRL-TK internal control (Promega) were generated using the Qiagen EndoFree Plasmid Maxi kit, and stored as purified DNA for future use.

Prior to transfection of cells, WT HMC3 cells were plated in 24-well plates in normal growth media and left to grow until reaching ∼80% confluence. Lipofectamine 3000:DNA complexes with P3000 reagent were generated in OptiMEM so that each well received 625ng plasmid DNA, 0.75μL of Lipofectamine 3000, and 1μL of P3000 reagent for a plasmid DNA (μg):Lipofectamine 3000 (μL) ratio of 1:1.2 per well. The vector DNA contained both the relevant experimental pGL4 plasmid, as well as the pRL-TK internal control plasmid at a ratio of 4.75:1. At 80% confluence, cells were given, in triplicate wells, the appropriate DNA:Lipofectamine 3000 complex, as well as mock transfected controls containing PBS in place of DNA, in their normal growth media, and allowed to incubate for approximately 48 hours before whole-cell lysates were harvested for luciferase assays using 70μL of ice-cold 1X Passive Lysis Buffer per well, according to manufacturer’s protocol (Promega).

Luciferase assays were carried out on whole cell lysates collected same day as assay using the Dual-Luciferase Assay® system according to manufacturer’s protocol (Promega). Luciferase Assay Reagent II and Stop&Glo® Reagent were prepared fresh on each day of assay. 20μL of each experimental condition lysate was plated in triplicate on a white, flat-bottom 96-well plate for a total of 9 wells/condition (3 wells of transfected cells per condition, each in triplicate). Each well was assay using the SpectraMax iD-5 plate reader by auto-injecting 100μL of Luciferase Assay Reagent II, waiting 2 seconds, measuring firefly luminescence for 10 seconds, injecting 100μL Stop&Glo Reagent, waiting 2 seconds, and measuring renilla luminescence for 10 seconds. Firefly:renilla ratios were calculated automatically by the plate reader.

### CRISPR-Cas9 Genome Editing in HMC3 Cells

CRISPR-Cas9 mediated deletion of the approximately 400bp region surrounding the proxy SNP (rs6024870) at the *CASS4* GWAS locus was achieved using a pooled lentiviral mCherry construct (lentiCRISPR v2-mCherry, Addgene 99154) containing one sgRNA on each side of the target region. sgRNAs were designed with CRISPOR^81^, purchased from Integrated DNA Technologies, and cloned into the lentiCRISPR v2-mCherry plasmid by modified Golden Gate reaction^82–84^. Accurate insertion of sgRNAs into the plasmid was checked via Sanger sequencing. 293T cells were plated in 10cm dishes and allowed to grow to 80% confluence before being transfected with equimolar ratios of both lentiCRISPR v2-mCherry sgRNA-containing constructs (or the vector alone without sgRNA targeting sequences for the mock transduced controls), packaging construct, and envelope construct. Transfection was achieved using Lipofectamine 2000 according to manufacturer’s protocol (ThermoFisher Scientific). Transfection media was replaced after 4.5 hours and replaced with normal growth media. Growth media containing lentivirus was collected after 24 hours, filtered through a 0.22μm filter, and stored at -80°C until infection of HMC3 cells.

Passage 6 HMC3 cells were plated on 10cm dishes and allowed to adhere for 24 hours in normal growth media. Growth media was aspirated and replaced with fresh growth media, appropriate filtered lentiviral pool, and polybrene (2μg/mL final concentration) to facilitate lentiviral transduction. Media was replaced after 48 hours, and mCherry+ transduced cells were selected by fluorescence-activated cell sorting (FACS) using the FACSJazz platform (BD Biosciences) at the CHOP FACS Core. mCherry+ stocks were both expanded to new 10cm dishes and frozen using normal growth media supplemented with 10% DMSO for storage in liquid nitrogen.

Single-cell CRISPR clones were obtained by sorting single mCherry+ HMC3 cells into individual wells of 96-well plates containing normal growth media by FACS using the MoFlo Astrios platform (Beckman Coulter) at the CHOP FACS Core. Clones were grown up before harvesting genomic DNA using the Quick DNA/RNA MiniPrep kit (Zymo) according to manufacturer’s protocol.

PCR primers flanking the targeting deletion region were used to amplify genomic DNA harvested from CRISPR-edited HMC3 clonal cells. PCR products for both mock transduced and knockout cells were the predicted sizes: mock transduced (WT) PCR product was 889bp, and the CRISPR deletion product was 495bp. Clones heterozygous for the deletion had both WT and deletion bands present on the PCR. To further confirm successful deletion, PCR products for HMC3 clones were subcloned into the TOPO-TA vector (Thermo Fisher), and positive clones were subsequently Sanger sequenced.

### RNA-Sequencing in HMC3 Cells

Total RNA was isolated from all cells by lifting cells with TRIzol reagent (Invitrogen) and whole RNA was isolated from lysates using the Direct-Zol RNA MiniPrep kit (Zymo) according to manufacturer’s protocol. Resulting RNA quality was assessed on an RNA 6000 Nano chip on the Agilent BioAnalyzer. rRNA was depleted using the QIAseq FastSelect RNA Removal kit (Qiagen). RNA-seq libraries were prepared using the NEBNext Ultra II Directional RNA Library Prep kit for Illumina (NEB) following manufacturer’s protocol.

### RNA-seq Analysis

The pair-end fastq files were mapped to genome assembly hg19 using STAR (v2.5.2b)^85^ independently for each replicate. GencodeV19 annotation was used for gene feature annotation and the raw read count for gene feature was calculated by htseq-count (v0.6.1)^86^ with parameter settings -f bam -r pos -s reverse -t exon -m union. The gene features localized on chrM or annotated as rRNAs were removed from the final sample-by-gene read count matrix.

The differential analysis was performed in R (v4.3.1) using R package edgeR (v3.42.4)^87^. Briefly, the raw reads on genes features were transformed in CPM (read Counts Per Million total reads). The gene features with less than 90 reads in any sample were removed from differential analysis. The trimmed mean of M-values (TMM) method were used to calculate normalization scaling factors and quasi-likelihood negative binomial generalized log-linear (glmQLFit) approach was applied to the count data with model fitting ∼0 + condition (i.e. KO or Mock). The differential expression genes (DEGs) between KO and mock cells were identified with cut-off FDR < 0.05 and absolute logFC > 0.58.

### Immunoblotting

Immunoblotting was performed using standard procedures in mock and CRISPR-edited homozygous knockout HMC3 cell clones with some modifications. 6-well plates of ∼90% confluent HMC3 cells were washed once with 2mL of ice-cold PBS per well, then protein was lysed and harvested in 1X RIPA lysis buffer (VWR Life Science) supplemented with 1X cOmplete^TM^ Mini EDTA-free protease inhibitor cocktail (Sigma). Cell lysates were collected and clarified with 15 minutes centrifugation at 14,000rpm at 4°C. The protein concentration of each lysate was determined using a BCA assay (Pierce). 3μg of each lysate was loaded into 4-12% bis-tris SDS-polyacrylamide gels (ThermoFisher), and electrotransferred onto polyvinyl difluoride membranes. In lieu of a loading control, we performed whole-protein staining for downstream quantification by incubating membranes for 25 minutes in No-Stain Protein Labeling Reagent (ThermoFisher Scientific), prepared according to manufacturer’s protocol. Membranes were blocked at room temperature for 1 hour in 5% non-fat skim milk in 1X TBST (tris-buffered saline containing 0.01% Tween-20), then incubated overnight at 4°C in primaries antibodies at dilutions recommended by the manufacturers (see below). Membranes were washed 3 times in 1X TBST, then incubated for 1 hour at room temperature with appropriate horseradish peroxidase-conjugated secondary antibodies. Finally, blots were incubated for 2 minutes in SuperSignal West Pico Chemilluminescent Substrate (Thermo Fisher) and imaged on the iBrightFL1000 (Thermo Fisher). Relative band intensities for proteins of interest were calculated using ImageJ^88^ following the protocol described by Hossein Davarinejad (https://www.yorku.ca/yisheng/Internal/Protocols/ImageJ.pdf). The following primary antibodies were used: RTFDC1 (Millipore-Sigma, HPA053986, 0.1μg/mL), CASS4 (Millipore Sigma, SAB1304501-400UL, 1:1000), DDK (FLAG) (Origene TA50011-100, 1:2000). The following secondary antibodies were used: Anti-rabbit IgG, HRP-linked antibody (Cell Signaling Technology, 7074S, 1:2500), Anti-mouse IgG, HRP-linked antibody (Cell Signaling Technology, 7076S, 1:2500).

### siRNA Depletion of HMC3 Cells

Wild type HMC3 cells were seeded into 6-well plates in normal media and growth conditions, and allowed to grow until reaching approximately 60% confluence. Transfection of cells with siRNA pools was achieved using the DharmaFECT 1 transfection reagent (Dharmacon Inc.) in a concentration of 5μL per well. To knock down *RTFDC1*, a pool of siRNAs targeting *RTFDC1* was utilized (Dharmacon L-020569-00-0010 ON-TARGETplus Human RTFDC1 (51507) siRNA SMARTpool), and a non-targeting control pool was used in tandem as a control (Dharmacon D-001810-10-05). siRNA pools were resuspended to 25μM stocks using 1X siRNA buffer (Dharmacon Inc.) and further diluted to 5μM in 1X siRNA buffer on day of transfection. HMC3 cells were transfected with 25μM siRNA pool per well according to manufacturer’s protocol, and allowed to grow at in serum- and antibiotic-free HMC3 growth media for 24 hours at normal growth conditions before transfection media was removed and replaced with fresh HMC3 growth media, and grown for an additional 48 hours. 72 hours total post-transfection, cells were harvested for downstream experiments, and knockdown efficiency was assessed by immunoblot.

### RTFDC1 Lentiviral Transduction for Overexpression

rs6024870 enhancer KO HMC3 clones were seeded in 6-well plates in normal media and growth conditions, and allowed to grow until reaching approximately 60% confluence. Cells were transduced with 40μL of either *RTFDC1-DDK* lentivirus (OriGene LentiOrf cat: RC201652L3V, >10^7^ TU/mL) or mock lentivirus (Origene LentiOrf cat: PS100092V, pLenti-C-Myc-DDK-P2A-Puro >1×10^7^ TU/mL) in normal HMC3 growth media supplemented with 2μg/mL of polybrene transduction reagent. Transductions were conducted in duplicate (2 wells at a time) to obtain 2 separate repliactes of each transduction. All transduced cells were grown in appropriate transduction cocktail for 24 hours at normal growth conditions before media was replaced with fresh HMC3 growth media and grown for an additional 24 hours. Transduced cells were then dosed with 2μg/mL puromycin and grown for 7 days to allow for selection of positively transduced cells. After puromycin selection, overexpression and correspondent mock lines were frozen for long-term storage, and cellular materials were harvested for downstream experiments. Successful overexpression of *RTFDC1* was assessed by immunoblotting for both endogenous RTFDC1, as well for DDK (OriGene TA50011-100, host: mouse, 1:2000 dilution).

### Cytokine ELISA

250,000 HMC3 cells of all experimental conditions were plated in 6-well plates, and allowed to grow at normal cell culture conditions until reaching 80% confluence. 24 hours prior to media collection, media was aspirated from 6-well plates containing 80% confluent HMC3 cells, and replaced with 2mL of fresh HMC3 growth media. 24 hours later, media was removed from individual wells and centrifuged for 10 minutes at 4°C at 1500rpm to remove cell debris. 1.5mL of supernatant was transferred to cryovials for storage at -80°C until ELISA was performed. ELISA for pro-inflammatory cytokines IL-6 and IL-8 were performed in technical triplicate according to manufacturer’s protocol (Thermo Fisher KHC0061 and Thermo Fisher KHC0081, respectively), and resulting assay plate was read for optical density on the ID5 SpectraMax plate reader at a 450nm wavelength. A new standard curve was performed on each plate. Standard curve and resulting sample concentrations were calculated by the plate reader.

### γH2AX Fluorescence Assay

The γH2AX fluorescence assays were performed in accordance to kit instructions with some modifications (AbCam ab242296). 24 hours prior to assay, 89,000 HMC3 cells were seeded in normal growth media into each individual well of 48-well cell culture treated plates. On the day of assay, growth media was aspirated from each cell well, and in non-treated controls, immediately replaced with fresh normal growth media. Cells for the hour 0 and hour 24 timepoints were treated with 200μL of normal growth media containing a 1:200 dilution of etoposide, as indicated by kit instructions. All cells were incubated for 1 hour at normal growth conditions before being fixed in 3.7% formaldehyde for 10 minutes. Cells were then washed with PBS before being permeabilized in 90% ice-cold methanol for 10 minutes, then washed again with PBS. Cells were then blocked for 30 minutes in blocking buffer (1% BSA in PBS), and then stained for γH2AX according to kit specifications. After the final wash step, cells were incubated in NucBlue nuclear stain (ThermoFisher Scientific) for 10 minutes, then washed with kit wash buffer (0.05% Tween in PBS) 5 times before adding 200μL DPBS for image acquisition. Each timepoint and genotype was plated in duplicate (2 wells) on two separate days, and each well imaged in 3 separate locations, for a total of 6 images per genotype/timepoint combination. Each image was acquired at 10X magnification for brightfield, DAPI, and FITC using an Olympus DP74 camera using appropriate fluorescent filters and Olympus cellSens Standard software. Corrected total cell fluorescence intensity (CTCFI) was then measured for the FITC fluorescence (converted to 8-bit) of each image using ImageJ^88^, circling each individual nucleus with the Freehand selection tool, utilizing the following calculation:

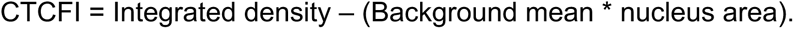

## ACKNOWLEDGEMENTS

E.A.B. is supported by the NIA (F31 AG074532). A.C. is supported by the NHGRI R35 HG011959. A.C. and S.F.A.G. are supported by the NIA (R01 AG057516). S.F.A.G. is also supported by the Daniel B. Burke Endowed Chair for Diabetes Research.

The results published here are in whole or in part based on data obtained from the AD Knowledge Portal (https://adknowledgeportal.org/). The data, analyses and tools are shared early in the research cycle without a publication embargo on secondary use. Data is available for general research use according to the following requirements for data access and data attribution. For access to content described in this manuscript see: http://doi.org/10.7303/syn26207321. Code used is available upon reasonable request from the corresponding authors. We thank the patients and families who donated material for these studies. We thank the computational resources and staff expertise provided by the Scientific Computing group at the Icahn School of Medicine at Mount Sinai.

The authors would like to thank Khanh Trang for assistance in generating the variant-to-gene mapping figure, Nicholas Wachowski and Winter Bruner for experimental advice and assistance, and Mitchell Conery, Mary Ann Weidekamp, and Max Dudek for assisting with data formatting and figure generation.

Some figures were generated with BioRender.

## AUTHOR CONTRIBUTIONS

E.A.B., S.F.A.G., C.D.B., and A.C. conceived the project. E.A.B., M.A., K.C., and S.H.L. performed cell culture. E.A.B tested cell culture for mycoplasma contamination. E.A.B. collected relevant cell materials. E.A.B. and M.A. prepared bulk RNA-seq libraries. S.L. and K.H. prepared bulk ATAC-seq libraries. M.A., J.A.P., and M.E.L. prepared Capture-C libraries. J.A.P. sequenced bulk RNA-seq, ATAC-seq, and Capture-C libraries. C.S., E.M., and A.C. processed and performed bioinformatic analyses, including bulk RNA-seq, ATAC-seq, and Capture-C analyses, including the variant-to-gene mapping from LOAD GWAS. E.A.B., M.E.J., and J.A.P. designed sgRNAs. E.A.B. performed CRISPR in HMC3 cells. E.A.B. and M.R. validated CRISPR clones for off-target effects. E.A.B. designed phenotypic validation experiments for CRISPR edited HMC3 cells. E.A.B. performed all ELISA assays and subsequent data analysis. E.A.B. and M.R. performed all γH2AX fluorescence assays. E.A.B. performed data analysis for γH2AX fluorescence assays. E.A.B. and S.H.L. designed luciferase constructs. E.A.B. optimized HMC3 cell transfection. E.A.B. and M.R. performed luciferase assays. E.A.B. performed luciferase assay data analysis. M.C.P. performed TF motif analysis. LS.W., G.D.S., J.A.P., A.D.W., S.A.A., S.F.A.G., and A.C. provided critical feedback and supervision. E.A.B., S.F.A.G., and A.C. wrote the original manuscript draft. All authors (except C.D.B) reviewed and edited the final manuscript.

## DATA AVAILABILITY

Upon publication of this manuscript all newly generated data sets will be available on the Gene Expression Omnibus (GEO).

## SUPPLEMENTARY FIGURES AND FIGURE LEGENDS

**Supplementary Figure 1:**
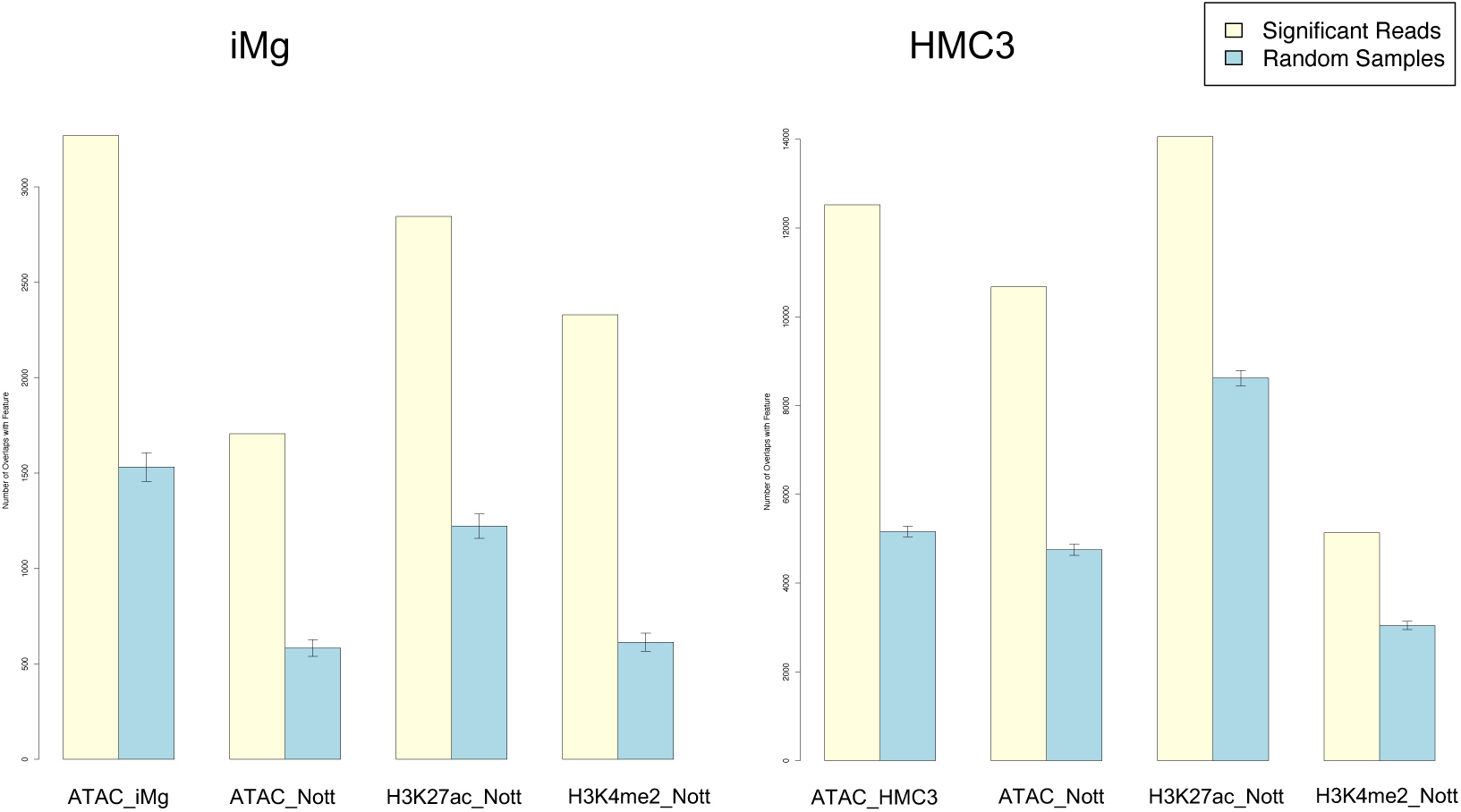
Promoter-interacting regions identified by our Capture-C experiments in iMg (left) and HMC3 (right) are enriched for active chromatin markers from human brain microglia. Yellow bars: number of overlaps with significant (CHiCAGO score>5) cis-interacting fragments (at 4-fragment resolution; bait-to-bait interactions were excluded); blue bars: expected overlaps based on 100 random subsets of fragments with a similar distribution of distances from the baits. ATAC-seq data is from our own experiments in iMg and HMC3 or Nott *et al*^22^. H3K27ac and H3K4me3 are from Nott *et al*^22^. Error bars represent 95% confidence intervals.

**Supplementary Figure 2:**
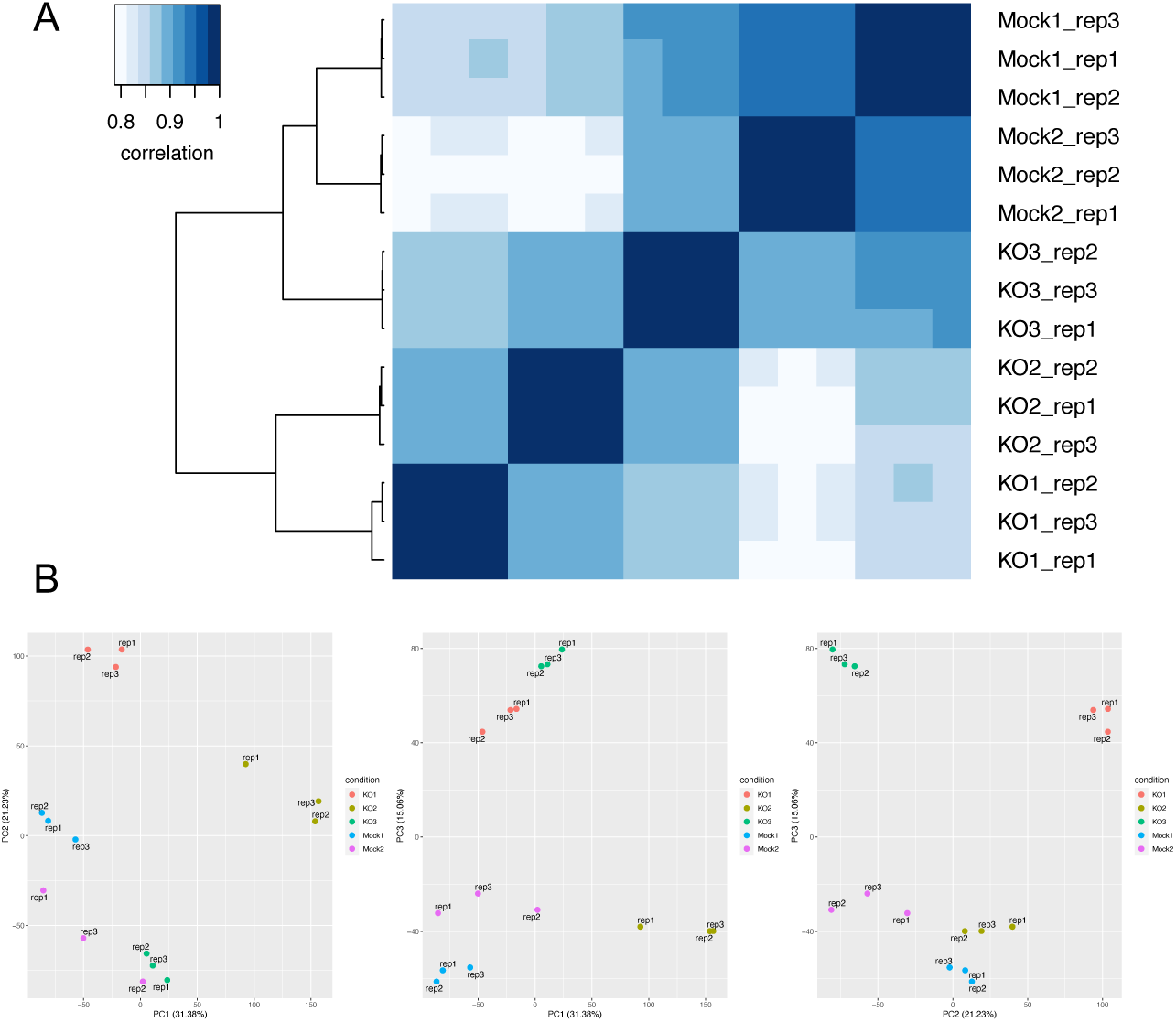
**(A)** Heatmap of the pairwise Pearson correlation of gene expression for 3 clones of CRISPR enhancer KO HMC3 cells and 2 mock controls, each with 3 technical replicates, clustered by distance. **(B)** Principal component analysis of the same datasets as in A.

**Supplementary Figure 3:**
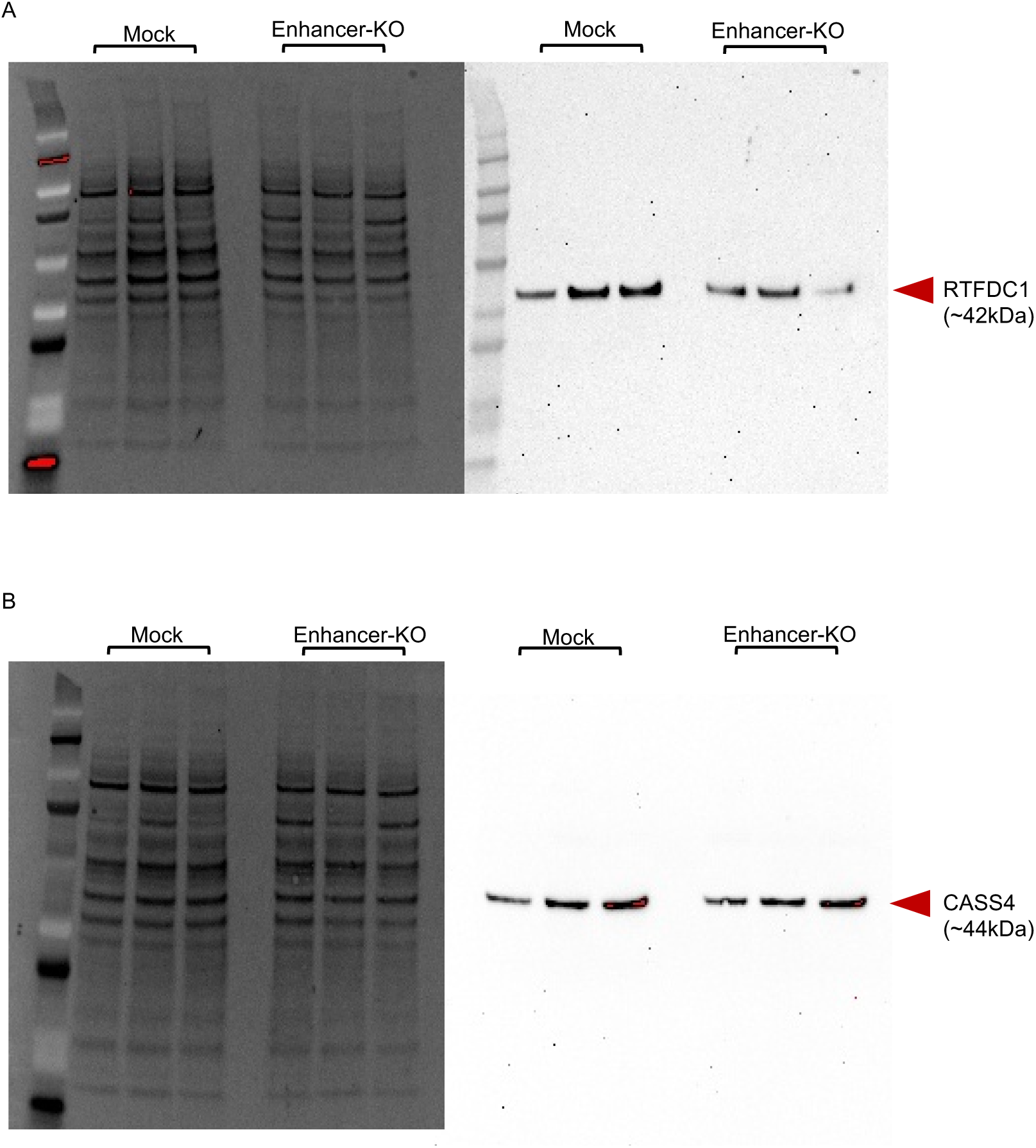
**(A)** Separated images of whole protein stain (left) and RTFDC1 (right) from immunoblot in Figure 3B. **(B)** Separated images of whole protein stain (left) and CASS4 (right) from immunoblot in Figure 3D.

**Supplementary Figure 4:**
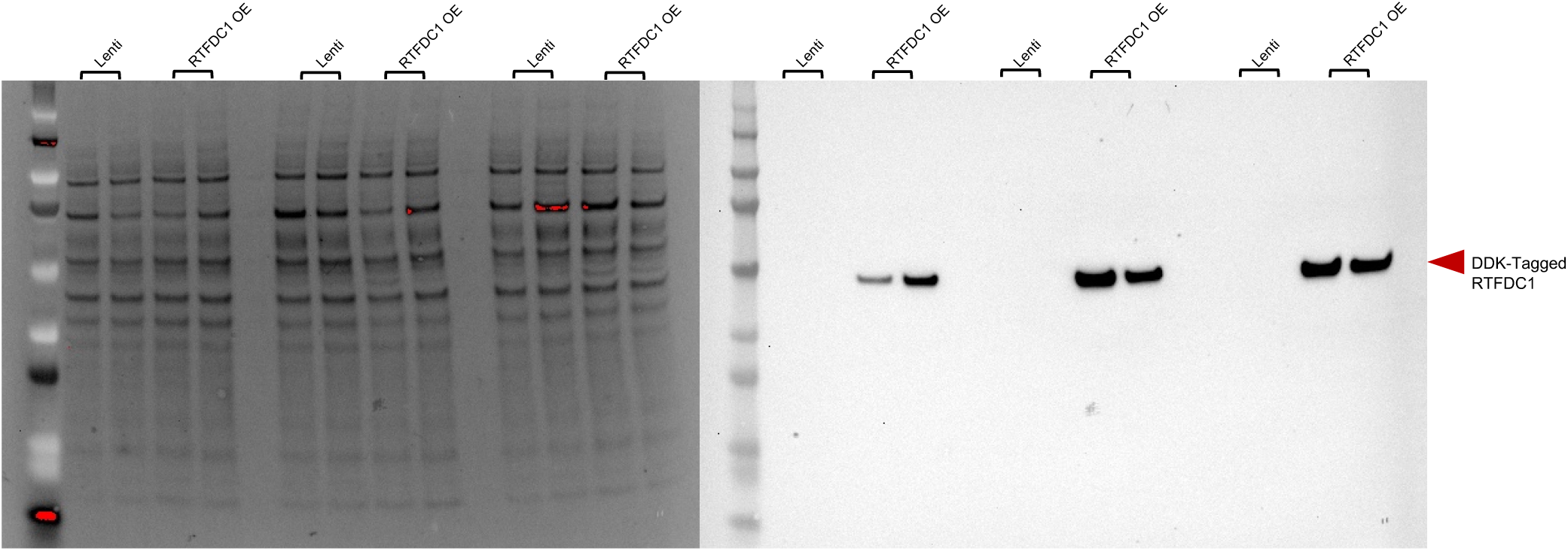
Separated images of whole protein stain (left) and DDK-tagged RTFDC1 (right) from immunoblot in Figure 4C (*RTFDC1* lentiviral overexpression system). Each lane represents a different lentiviral transduction.

**Supplementary Figure 5:**
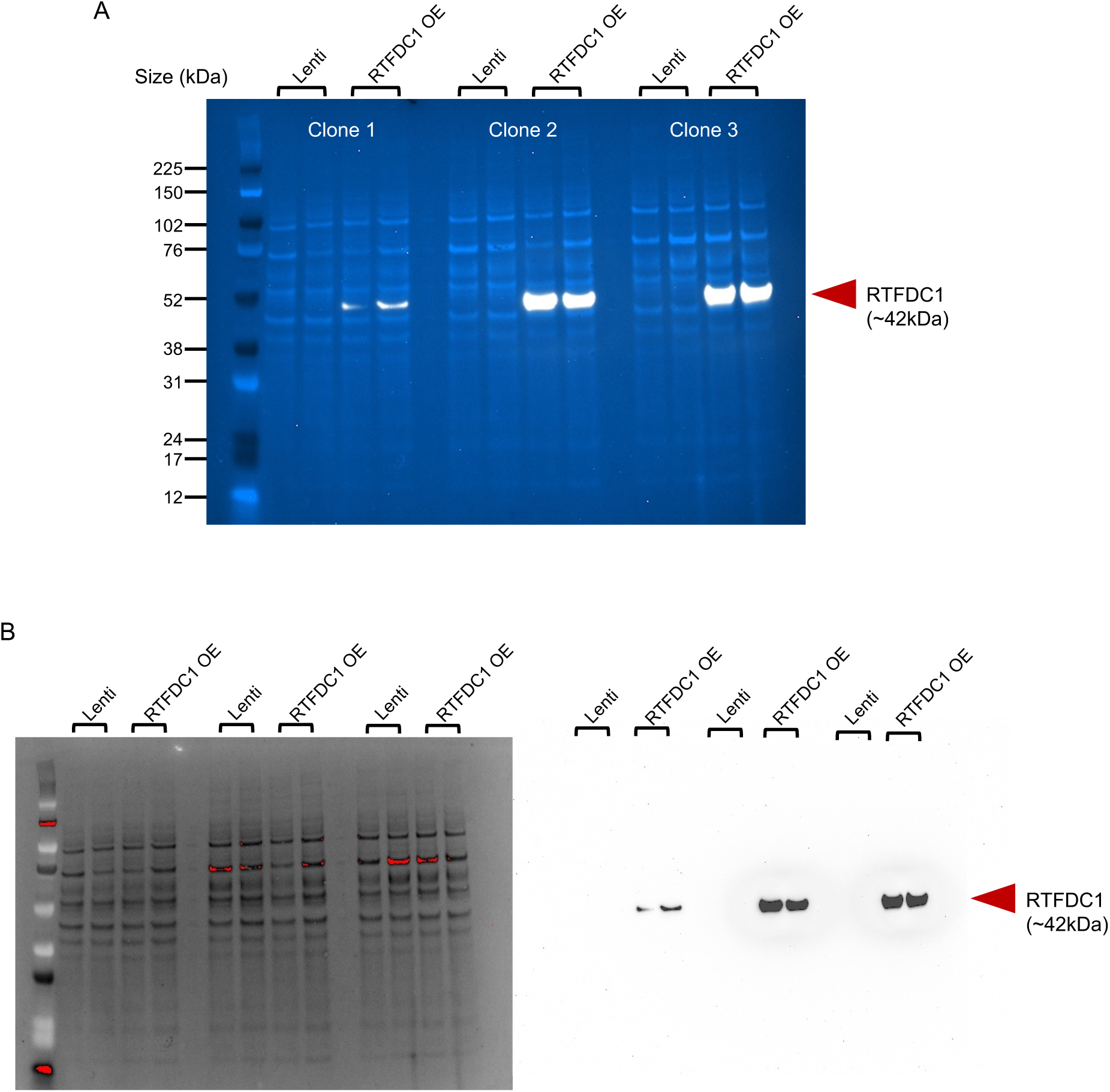
**(A)** Immunoblot for endogenous RTFDC1 from the same samples as Figure 4C. Lenti: mock transduced. RTFDC1 OE: transduced with DDK-tagged *RTFDC1* ORF. White band: DDK-tagged RTFDC1. Light blue bands: whole protein stain. **(B)** Separated images of whole protein stain (left) and RTFDC1 (right) from A.

**Supplementary Figure 6:**
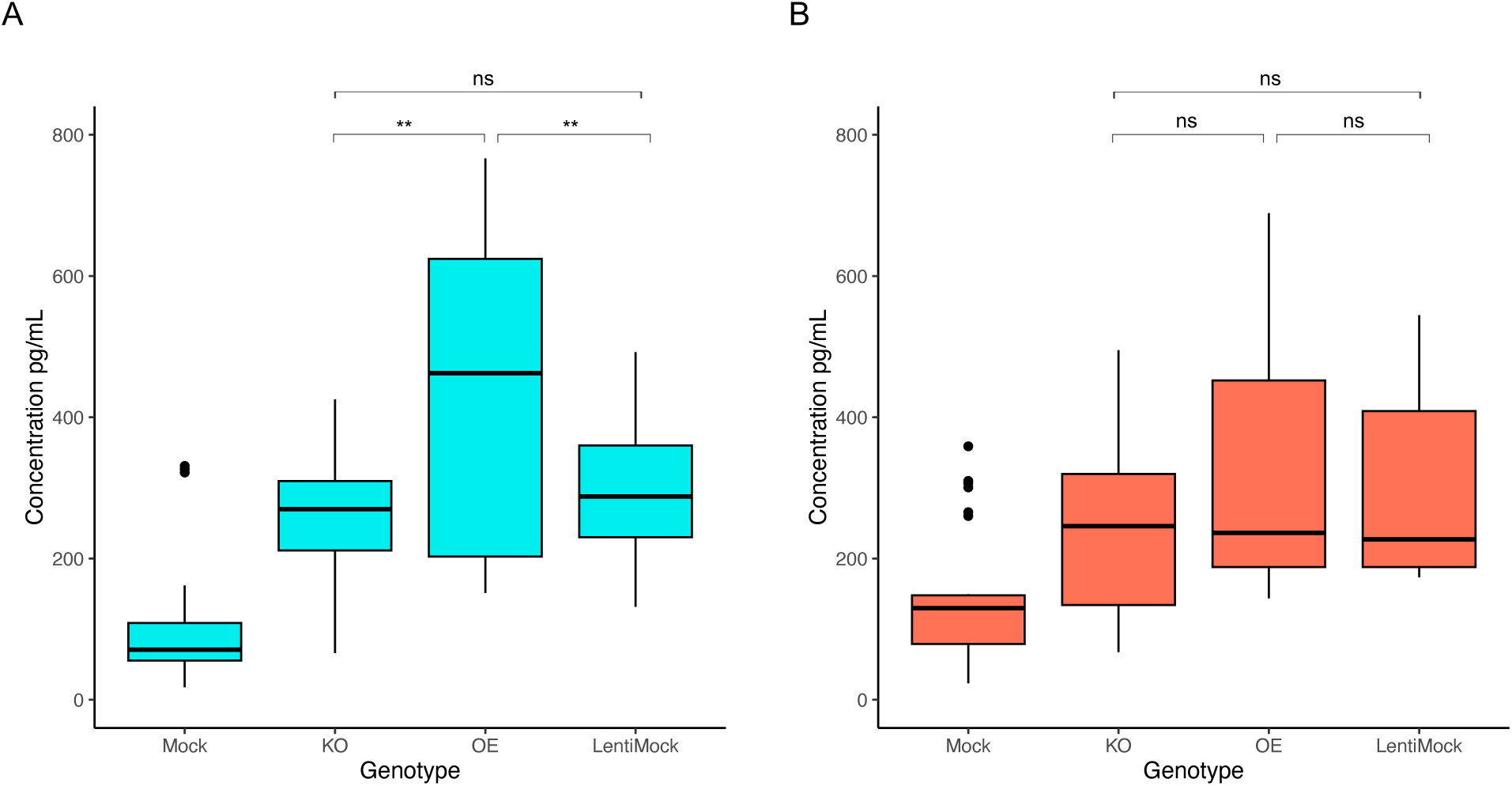
**(A)** Same as Figure 4D for IL-8 secretion, but including mock lentiviral overexpression vector (LentiMock) in the analysis. ** *P*<0.001 by one-way ANOVA with Tukey’s HSD. **(B)** Same as Figure 4E for IL-6 secretion, but including mock lentiviral overexpression vector (LentiMock) in the analysis. One-way ANOVA with Tukey’s HSD.

**Supplementary Figure 7:**
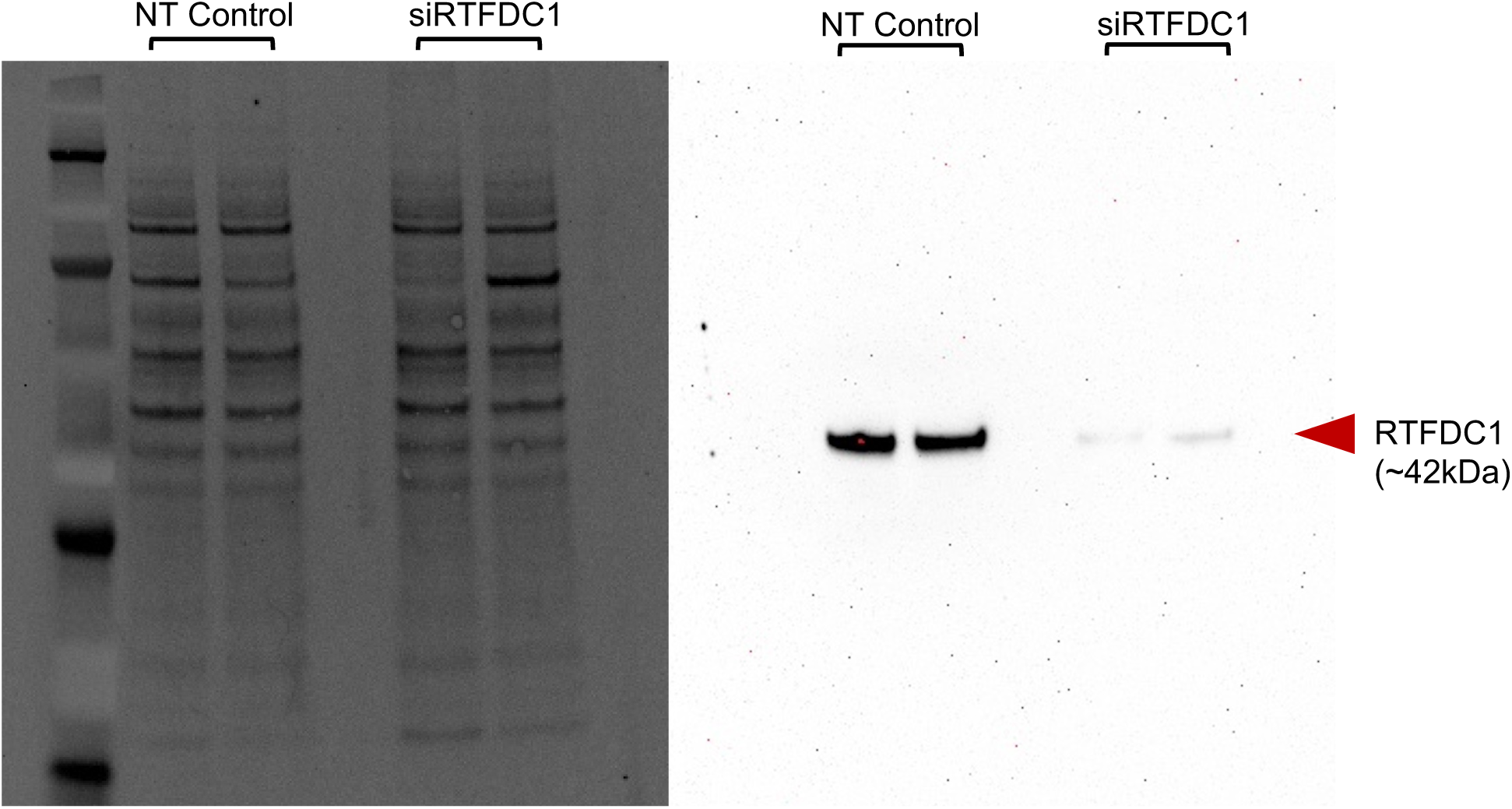
Separated images of whole protein stain (left) and RTFDC1 (right) from immunoblot in Figure 5C (siRNA knockdown of RTFDC1). Each lane represents an individual siRNA transfection.

**Supplementary Figure 8:**
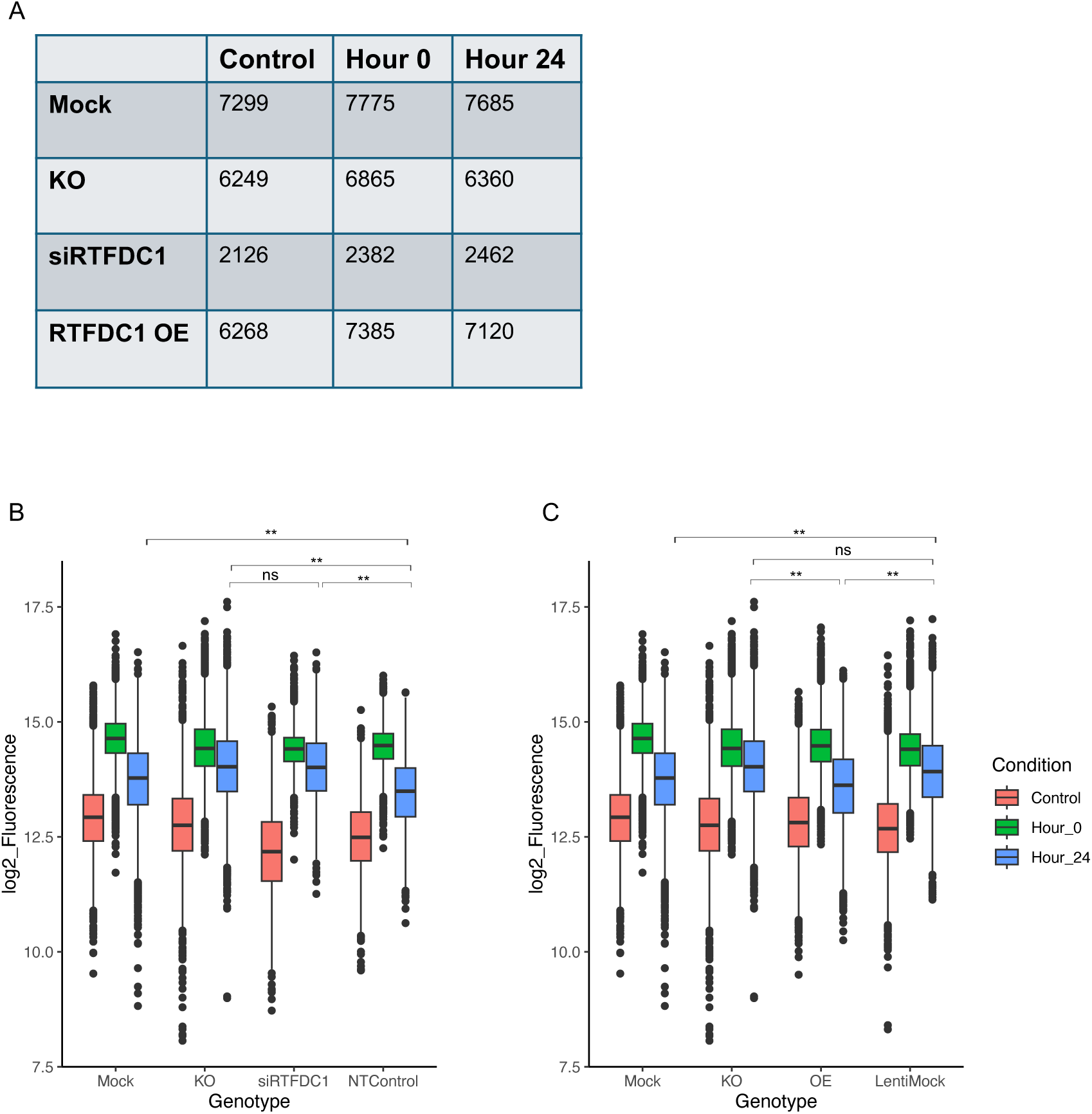
**(A)** Table displaying number of fluorescent nuclei quantified for each genotype and condition in the γH2AX staining experiment. **(B)** Same as Figure 5D, but including non-targeting siRNA control (NTControl) in the analysis. ** *P*<2.2×10^−16^ by two-way ANOVA (genotype by condition) with Tukey’s HSD. **(C)** Same as Figure 5E, but including mock lentiviral overexpression vector (LentiMock) in the analysis. ** *P*<2.2×10^−16^ by two-way ANOVA with Tukey’s HSD.

**Supplementary Figure 9:**
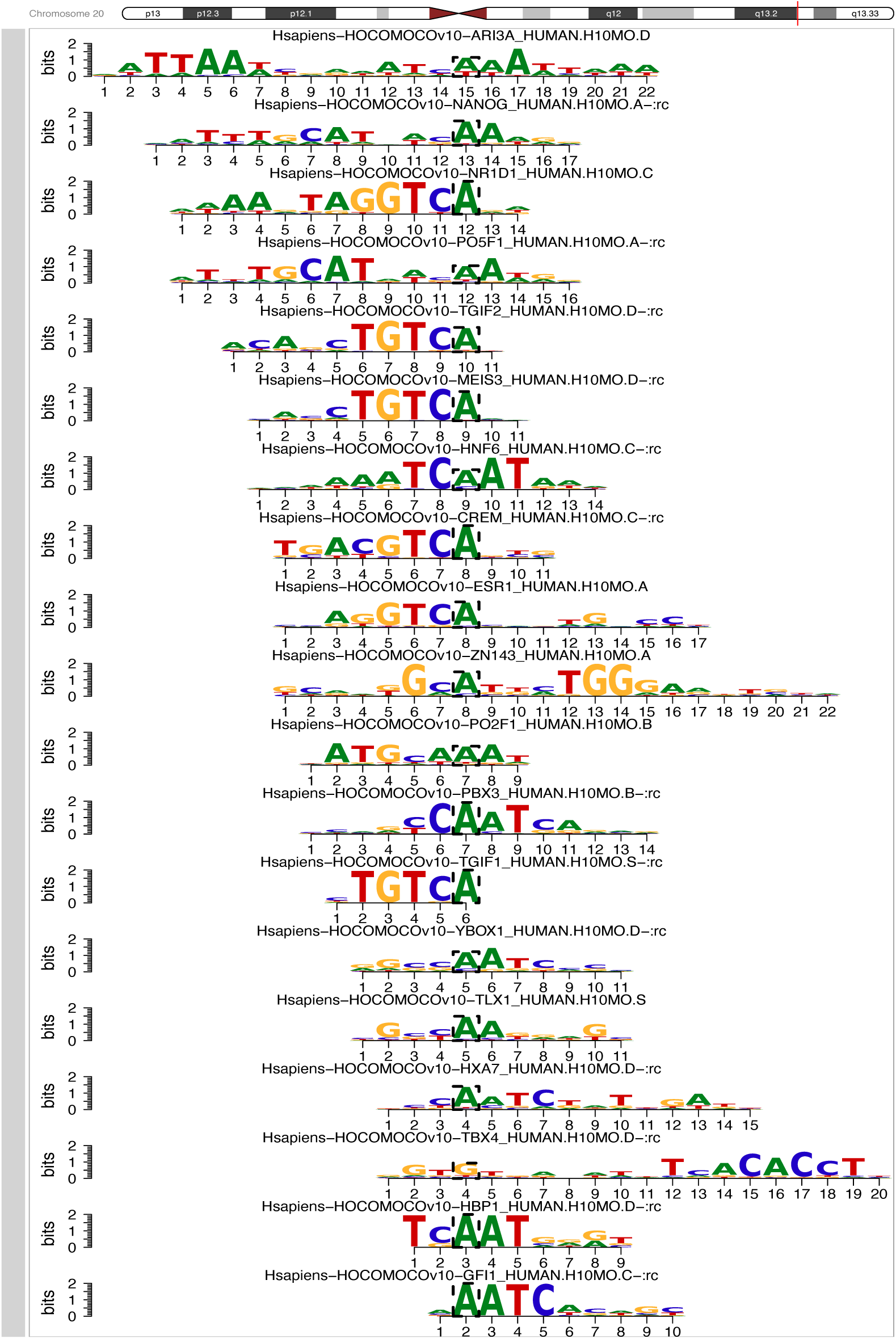
Complete list of transcription factor motifs with differential binding affinity relative to the rs6024870 protective A allele. Figure generated with MotifBreakR.

## REFERENCES

1 Alzheimer’s Disease Facts and Figures. (2023).

2 Reitz, C., Brayne, C. & Mayeux, R. Epidemiology of Alzheimer disease. Nat Rev Neurol 7, 137–152 (2011). 10.1038/nrneurol.2011.2

3 Gatz, M. et al. Role of genes and environments for explaining Alzheimer disease. Arch Gen Psychiatry 63, 168–174 (2006). 10.1001/archpsyc.63.2.168

4 Bellenguez, C. et al. New insights into the genetic etiology of Alzheimer’s disease and related dementias. Nat Genet 54, 412–436 (2022). 10.1038/s41588-022-01024-z

5 Jansen, I. E. et al. Genome-wide meta-analysis identifies new loci and functional pathways influencing Alzheimer’s disease risk. Nat Genet 51, 404–413 (2019). 10.1038/s41588-018-0311-9

6 El Khoury, J., et al. Scavenger receptor-mediated adhesion of microglia to beta-amyloid fibrils. Nature 382, 716–719 (1996). 10.1038/382716a0

7 Frenkel, D. et al. Scara1 deficiency impairs clearance of soluble amyloid-beta by mononuclear phagocytes and accelerates Alzheimer’s-like disease progression. Nat Commun 4, 2030 (2013). 10.1038/ncomms3030

8 El Khoury, J. B., et al. CD36 mediates the innate host response to beta-amyloid. J Exp Med 197, 1657–1666 (2003). 10.1084/jem.20021546

9 Gold, M. & El Khoury, J. beta-amyloid, microglia, and the inflammasome in Alzheimer’s disease. Semin Immunopathol 37, 607–611 (2015). 10.1007/s00281-015-0518-0

10 Sarlus, H. & Heneka, M. T. Microglia in Alzheimer’s disease. J Clin Invest 127, 3240–3249 (2017). 10.1172/JCI90606

11 Venegas, C. et al. Microglia-derived ASC specks cross-seed amyloid-beta in Alzheimer’s disease. Nature 552, 355–361 (2017). 10.1038/nature25158

12 Tansey, K. E., Cameron, D. & Hill, M. J. Genetic risk for Alzheimer’s disease is concentrated in specific macrophage and microglial transcriptional networks. Genome Med 10, 14 (2018). 10.1186/s13073-018-0523-8

13 Sims, R. et al. Rare coding variants in PLCG2, ABI3, and TREM2 implicate microglial-mediated innate immunity in Alzheimer’s disease. Nat Genet 49, 1373–1384 (2017). 10.1038/ng.3916

14 Jin, S. C. et al. Coding variants in TREM2 increase risk for Alzheimer’s disease. Hum Mol Genet 23, 5838–5846 (2014). 10.1093/hmg/ddu277

15 Altshuler, D., Daly, M. J. & Lander, E. S. Genetic mapping in human disease. Science 322, 881–888 (2008). 10.1126/science.1156409

16 Corradin, O. & Scacheri, P. C. Enhancer variants: evaluating functions in common disease. Genome Med 6, 85 (2014). 10.1186/s13073-014-0085-3

17 Smemo, S. et al. Obesity-associated variants within FTO form long-range functional connections with IRX3. Nature 507, 371–375 (2014). 10.1038/nature13138

18 Claussnitzer, M. et al. FTO Obesity Variant Circuitry and Adipocyte Browning in Humans. N Engl J Med 373, 895–907 (2015). 10.1056/NEJMoa1502214

19 Rosen, C. J. & Ingelfinger, J. R. Unraveling the Function of FTO Variants. N Engl J Med 373, 964–965 (2015). 10.1056/NEJMe1508683

20 Chesi, A. et al. Genome-scale Capture C promoter interactions implicate effector genes at GWAS loci for bone mineral density. Nat Commun 10, 1260 (2019). 10.1038/s41467-019-09302-x

21 Andersson, R. et al. An atlas of active enhancers across human cell types and tissues. Nature 507, 455–461 (2014). 10.1038/nature12787

22 Nott, A. et al. Brain cell type-specific enhancer-promoter interactome maps and disease-risk association. Science 366, 1134–1139 (2019). 10.1126/science.aay0793

23 Kunkle, B. W. et al. Genetic meta-analysis of diagnosed Alzheimer’s disease identifies new risk loci and implicates Abeta, tau, immunity and lipid processing. Nat Genet 51, 414–430 (2019). 10.1038/s41588-019-0358-2

24 Pahl, M. C. et al. Cis-regulatory architecture of human ESC-derived hypothalamic neuron differentiation aids in variant-to-gene mapping of relevant complex traits. Nat Commun 12, 6749 (2021). 10.1038/s41467-021-27001-4

25 Su, C. et al. Mapping effector genes at lupus GWAS loci using promoter Capture-C in follicular helper T cells. Nat Commun 11, 3294 (2020). 10.1038/s41467-020-17089-5

26 Su, C. et al. 3D promoter architecture re-organization during iPSC-derived neuronal cell differentiation implicates target genes for neurodevelopmental disorders. Prog Neurobiol 201, 102000 (2021). 10.1016/j.pneurobio.2021.102000

27 Lasconi, C. et al. Variant-to-Gene-Mapping Analyses Reveal a Role for the Hypothalamus in Genetic Susceptibility to Inflammatory Bowel Disease. Cell Mol Gastroenterol Hepatol 11, 667–682 (2021). 10.1016/j.jcmgh.2020.10.004

28 Su, C. et al. 3D chromatin maps of the human pancreas reveal lineage-specific regulatory architecture of T2D risk. Cell Metab 34, 1394–1409 e1394 (2022). 10.1016/j.cmet.2022.08.014

29 Cousminer, D. L. et al. Genome-wide association study implicates novel loci and reveals candidate effector genes for longitudinal pediatric bone accrual. Genome Biol 22, 1 (2021). 10.1186/s13059-020-02207-9

30 Littleton, S. H. et al. Variant-to-function analysis of the childhood obesity chr12q13 locus implicates rs7132908 as a causal variant within the 3’ UTR of FAIM2. Cell Genom 4, 100556 (2024). 10.1016/j.xgen.2024.100556

31 Wightman, D. P. et al. A genome-wide association study with 1,126,563 individuals identifies new risk loci for Alzheimer’s disease. Nat Genet 53, 1276–1282 (2021). 10.1038/s41588-021-00921-z

32 Arnold, M., Raffler, J., Pfeufer, A., Suhre, K. & Kastenmuller, G. SNiPA: an interactive, genetic variant-centered annotation browser. *Bioinformatics (Oxford*, England) 31, 1334–1336 (2015). 10.1093/bioinformatics/btu779

33 Ryan, S. K. et al. Neuroinflammation and EIF2 Signaling Persist despite Antiretroviral Treatment in an hiPSC Tri-culture Model of HIV Infection. Stem Cell Reports 14, 703–716 (2020). 10.1016/j.stemcr.2020.02.010

34 Li, B. et al. NOX4 expression in human microglia leads to constitutive generation of reactive oxygen species and to constitutive IL-6 expression. J Innate Immun 1, 570–581 (2009). 10.1159/000235563

35 Medvinsky, A., Rybtsov, S. & Taoudi, S. Embryonic origin of the adult hematopoietic system: advances and questions. Development 138, 1017–1031 (2011). 10.1242/dev.040998

36 Ajami, B., Bennett, J. L., Krieger, C., Tetzlaff, W. & Rossi, F. M. Local self-renewal can sustain CNS microglia maintenance and function throughout adult life. Nat Neurosci 10, 1538–1543 (2007). 10.1038/nn2014

37 Prinz, M., Erny, D. & Hagemeyer, N. Ontogeny and homeostasis of CNS myeloid cells. Nat Immunol 18, 385–392 (2017). 10.1038/ni.3703

38 Ginhoux, F. et al. Fate mapping analysis reveals that adult microglia derive from primitive macrophages. Science 330, 841–845 (2010). 10.1126/science.1194637

39 Kierdorf, K. et al. Microglia emerge from erythromyeloid precursors via Pu.1- and Irf8-dependent pathways. Nat Neurosci 16, 273–280 (2013). 10.1038/nn.3318

40 Kosoy, R. et al. Genetics of the human microglia regulome refines Alzheimer’s disease risk loci. Nat Genet 54, 1145–1154 (2022). 10.1038/s41588-022-01149-1

41 Consortium, G. T. The GTEx Consortium atlas of genetic regulatory effects across human tissues. Science 369, 1318–1330 (2020). 10.1126/science.aaz1776

42 Song, R. et al. IRF1 governs the differential interferon-stimulated gene responses in human monocytes and macrophages by regulating chromatin accessibility. Cell Rep 34, 108891 (2021). 10.1016/j.celrep.2021.108891

43 Pahl, M. C. et al. Implicating effector genes at COVID-19 GWAS loci using promoter-focused Capture-C in disease-relevant immune cell types. Genome Biol 23, 125 (2022). 10.1186/s13059-022-02691-1

44 Wingett, S. et al. HiCUP: pipeline for mapping and processing Hi-C data. F1000Res 4, 1310 (2015). 10.12688/f1000research.7334.1

45. Cairns, J., et al. CHiCAGO: robust detection of DNA looping interactions in Capture Hi-C data. Genome Biol 17, 127 (2016). 10.1186/s13059-016-0992-2

46 Su, C., Pahl, M. C., Grant, S. F. A. & Wells, A. D. Restriction enzyme selection dictates detection range sensitivity in chromatin conformation capture-based variant-to-gene mapping approaches. Hum Genet 140, 1441–1448 (2021). 10.1007/s00439-021-02326-8

47 Finucane, H. K. et al. Partitioning heritability by functional annotation using genome-wide association summary statistics. Nat Genet 47, 1228–1235 (2015). 10.1038/ng.3404

48 Schizophrenia Working Group of the Psychiatric Genomics, C. Biological insights from 108 schizophrenia-associated genetic loci. Nature 511, 421–427 (2014). 10.1038/nature13595

49 Consortium, E. P. et al. Expanded encyclopaedias of DNA elements in the human and mouse genomes. Nature 583, 699–710 (2020). 10.1038/s41586-020-2493-4

50 Wang, X. et al. Genetic determinants of disease progression in Alzheimer’s disease. J Alzheimers Dis 43, 649–655 (2015). 10.3233/JAD-140729

51 Dourlen, P. et al. Functional screening of Alzheimer risk loci identifies PTK2B as an in vivo modulator and early marker of Tau pathology. Mol Psychiatry 22, 874–883 (2017). 10.1038/mp.2016.59

52 Mootha, V. K. et al. PGC-1alpha-responsive genes involved in oxidative phosphorylation are coordinately downregulated in human diabetes. Nat Genet 34, 267–273 (2003). 10.1038/ng1180

53 Subramanian, A. et al. Gene set enrichment analysis: a knowledge-based approach for interpreting genome-wide expression profiles. Proc Natl Acad Sci U S A 102, 15545–15550 (2005). 10.1073/pnas.0506580102

54 Kalman, J. et al. Serum interleukin-6 levels correlate with the severity of dementia in Down syndrome and in Alzheimer’s disease. Acta Neurol Scand 96, 236–240 (1997). 10.1111/j.1600-0404.1997.tb00275.x

55 Alsadany, M. A., Shehata, H. H., Mohamad, M. I. & Mahfouz, R. G. Histone deacetylases enzyme, copper, and IL-8 levels in patients with Alzheimer’s disease. Am J Alzheimers Dis Other Demen 28, 54–61 (2013). 10.1177/1533317512467680

56 Zhang, J. et al. CSF multianalyte profile distinguishes Alzheimer and Parkinson diseases. Am J Clin Pathol 129, 526–529 (2008). 10.1309/W01Y0B808EMEH12L

57 Swardfager, W. et al. A meta-analysis of cytokines in Alzheimer’s disease. Biol Psychiatry 68, 930–941 (2010). 10.1016/j.biopsych.2010.06.012

58 Hjorth, E., Frenkel, D., Weiner, H. & Schultzberg, M. Effects of immunomodulatory substances on phagocytosis of abeta(1-42) by human microglia. Int J Alzheimers Dis 2010 (2010). 10.4061/2010/798424

59 Rajalakshmy, A. R., Malathi, J. & Madhavan, H. N. Hepatitis C Virus NS3 Mediated Microglial Inflammation via TLR2/TLR6 MyD88/NF-kappaB Pathway and Toll Like Receptor Ligand Treatment Furnished Immune Tolerance. PLoS One 10, e0125419 (2015). 10.1371/journal.pone.0125419

60 Huang, X., Tanaka, T., Kurose, A., Traganos, F. & Darzynkiewicz, Z. Constitutive histone H2AX phosphorylation on Ser-139 in cells untreated by genotoxic agents is cell-cycle phase specific and attenuated by scavenging reactive oxygen species. Int J Oncol 29, 495–501 (2006).

61 Spitz, F. & Furlong, E. E. Transcription factors: from enhancer binding to developmental control. Nat Rev Genet 13, 613–626 (2012). 10.1038/nrg3207

62 Melhuish, T. A., Gallo, C. M. & Wotton, D. TGIF2 interacts with histone deacetylase 1 and represses transcription. J Biol Chem 276, 32109–32114 (2001). 10.1074/jbc.M103377200

63 Bollaert, E. et al. HBP1 phosphorylation by AKT regulates its transcriptional activity and glioblastoma cell proliferation. Cell Signal 44, 158–170 (2018). 10.1016/j.cellsig.2018.01.014

64 Cho, H. et al. Regulation of circadian behaviour and metabolism by REV-ERB-alpha and REV-ERB-beta. Nature 485, 123–127 (2012). 10.1038/nature11048

65 Yang, X. et al. Functional characterization of Alzheimer’s disease genetic variants in microglia. Nat Genet 55, 1735–1744 (2023). 10.1038/s41588-023-01506-8

66 Hughes, J. R. et al. Analysis of hundreds of cis-regulatory landscapes at high resolution in a single, high-throughput experiment. Nat Genet 46, 205–212 (2014). 10.1038/ng.2871

67 Fang, R. et al. Mapping of long-range chromatin interactions by proximity ligation-assisted ChIP-seq. Cell Res 26, 1345–1348 (2016). 10.1038/cr.2016.137

68 Mostafavi, H., Spence, J. P., Naqvi, S. & Pritchard, J. K. Systematic differences in discovery of genetic effects on gene expression and complex traits. Nature genetics 55, 1866–1875 (2023). 10.1038/s41588-023-01529-1

69 Kottemann, M. C., Conti, B. A., Lach, F. P. & Smogorzewska, A. Removal of RTF2 from Stalled Replisomes Promotes Maintenance of Genome Integrity. Mol Cell 69, 24–35 e25 (2018). 10.1016/j.molcel.2017.11.035

70 Conti, B. A. et al. RTF2 controls replication repriming and ribonucleotide excision at the replisome. Nat Commun 15, 1943 (2024). 10.1038/s41467-024-45947-z

71 Wingo, A. P. et al. Integrating human brain proteomes with genome-wide association data implicates new proteins in Alzheimer’s disease pathogenesis. Nat Genet 53, 143–146 (2021). 10.1038/s41588-020-00773-z

72 Nasser, J. et al. Genome-wide enhancer maps link risk variants to disease genes. Nature 593, 238–243 (2021). 10.1038/s41586-021-03446-x

73 Fulco, C. P. et al. Activity-by-contact model of enhancer-promoter regulation from thousands of CRISPR perturbations. Nat Genet 51, 1664–1669 (2019). 10.1038/s41588-019-0538-0

74 Phanstiel, D. H. et al. Static and Dynamic DNA Loops form AP-1-Bound Activation Hubs during Macrophage Development. Mol Cell 67, 1037–1048 e1036 (2017). 10.1016/j.molcel.2017.08.006

75 Huang, Y. & Mahley, R. W. Apolipoprotein E: structure and function in lipid metabolism, neurobiology, and Alzheimer’s diseases. Neurobiol Dis 72 **Pt A**, 3-12 (2014). 10.1016/j.nbd.2014.08.025

76 Martinez-Martinez, A. B. et al. Beyond the CNS: The many peripheral roles of APOE. Neurobiol Dis 138, 104809 (2020). 10.1016/j.nbd.2020.104809

77 Ostendorf, B. N. et al. Common germline variants of the human APOE gene modulate melanoma progression and survival. Nat Med 26, 1048–1053 (2020). 10.1038/s41591-020-0879-3

78 Janabi, N., Peudenier, S., Heron, B., Ng, K. H. & Tardieu, M. Establishment of human microglial cell lines after transfection of primary cultures of embryonic microglial cells with the SV40 large T antigen. Neurosci Lett 195, 105–108 (1995). 10.1016/0304-3940(94)11792-h

79 Maguire, J. A. et al. Generation of human control iPS cell line CHOPWT10 from healthy adult peripheral blood mononuclear cells. Stem Cell Res 16, 338–341 (2016). 10.1016/j.scr.2016.02.017

80 Paluru, P. et al. The negative impact of Wnt signaling on megakaryocyte and primitive erythroid progenitors derived from human embryonic stem cells. Stem Cell Res 12, 441–451 (2014). 10.1016/j.scr.2013.12.003

81 Concordet, J. P. & Haeussler, M. CRISPOR: intuitive guide selection for CRISPR/Cas9 genome editing experiments and screens. Nucleic Acids Res 46, W242–W245 (2018). 10.1093/nar/gky354

82 Engler, C., Kandzia, R. & Marillonnet, S. A one pot, one step, precision cloning method with high throughput capability. PLoS One 3, e3647 (2008). 10.1371/journal.pone.0003647

83 Engler, C., Gruetzner, R., Kandzia, R. & Marillonnet, S. Golden gate shuffling: a one-pot DNA shuffling method based on type IIs restriction enzymes. PLoS One 4, e5553 (2009). 10.1371/journal.pone.0005553

84 Engler, C. & Marillonnet, S. Generation of families of construct variants using golden gate shuffling. Methods Mol Biol 729, 167–181 (2011). 10.1007/978-1-61779-065-2_11

85 Dobin, A. et al. STAR: ultrafast universal RNA-seq aligner. Bioinformatics 29, 15–21 (2013). 10.1093/bioinformatics/bts635

86 Anders, S., Pyl, P. T. & Huber, W. HTSeq--a Python framework to work with high-throughput sequencing data. Bioinformatics 31, 166–169 (2015). 10.1093/bioinformatics/btu638

87 Robinson, M. D., McCarthy, D. J. & Smyth, G. K. edgeR: a Bioconductor package for differential expression analysis of digital gene expression data. Bioinformatics 26, 139–140 (2010). 10.1093/bioinformatics/btp616

88 Schneider, C. A., Rasband, W. S. & Eliceiri, K. W. NIH Image to ImageJ: 25 years of image analysis. Nat Methods 9, 671–675 (2012). 10.1038/nmeth.2089

89 Shi, L. et al. IL-1 Transcriptional Responses to Lipopolysaccharides Are Regulated by a Complex of RNA Binding Proteins. J Immunol 204, 1334–1344 (2020). 10.4049/jimmunol.1900650

